# Combining evolutionary and assay-labelled data for protein fitness prediction

**DOI:** 10.1101/2021.03.28.437402

**Authors:** Chloe Hsu, Hunter Nisonoff, Clara Fannjiang, Jennifer Listgarten

**Author notes:** Correspondence to: Chloe Hsu and Jennifer Listgarten.

## Abstract

Predictive modelling of protein properties has become increasingly important to the field of machine-learning guided protein engineering. In one of the two existing approaches, evolutionarily-related sequences to a query protein drive the modelling process, without any property measurements from the laboratory. In the other, a set of protein variants of interest are assayed, and then a supervised regression model is estimated with the assay-labelled data. Although a handful of recent methods have shown promise in combining the evolutionary and supervised approaches, this hybrid problem has not been examined in depth, leaving it unclear how practitioners should proceed, and how method developers should build on existing work. Herein, we present a systematic assessment of methods for protein fitness prediction when evolutionary and assay-labelled data are available. We find that a simple baseline approach we introduce is competitive with and often outperforms more sophisticated methods. Moreover, our simple baseline is plug-and-play with a wide variety of established methods, and does not add any substantial computational burden. Our analysis highlights the importance of systematic evaluations and sufficient baselines.

---

Naturally occurring proteins serve many crucial functions in maintaining life, but have also been co-opted for human endeavors such as gene editing^1,2^; lighting up specific parts of cells^3^; therapeutic drugs^4^; and herbicide-resistant crops^5^. Furthermore, in many cases we re-engineer proteins to better serve our needs. For example, we might enhance the original function, such as when we increase enzyme activity^6^ or make green fluorescent proteins (GFPs) brighter^7^. Alternatively, we might modify the original function to a related but different one, such as when we change an antibody to bind to a new target^8^. The two most common approaches to protein engineering are the iterative, laboratory-based method of directed evolution^9^, and the computational methods for physics-based rational design^10,11^.

In recent years, efforts have been made to use machine learning (ML) for improved protein engineering, toward the goal of achieving more desirable engineered proteins, or getting there with fewer laboratory experiments, or both. When the design space—the part of the protein being engineered—is constrained to be relatively small (*e. g.*, four amino acid positions^12^), an ML-based protein fitness model can be used to systematically screen all protein variants in the design space^12–15^. When the space becomes too large, a layer of optimization must be added because a comprehensive screen is no longer possible^16–19^. In both cases, a key component of the ML-guided protein engineering approach is a reliance on an accurate ML-based fitness model—one that predicts protein property from protein sequence^20–25^.

There have been two main strategies for estimating protein fitness models. The first strategy leverages implicit fitness constraints present in naturally occurring protein sequences, so-called *evolutionary data*. Evolutionary approaches start from a single query protein with the presence of property of interest (*e. g.*, a particular GFP that fluoresces), and search through databases of naturally occurring proteins to find a set of related proteins—typically by sequence homology—that are assumed to be enriched for the same property as the query sequence. Then, a density model of this set of protein sequences is estimated, and the sequence density is used as a proxy for predicting fitness rankings^20,21^. Over the years, a set of increasingly richer density models has been employed for this purpose, ranging from position-specific substitution matrices^28^, profile Hidden Markov Models (HMMs)^29^, Potts models^14,20^, to Variational Autoencoders (VAEs)^21^ and long short-term memory networks (LSTMs)^30^. Some of these models assume that the evolutionarily-related proteins have already been aligned into a Multiple Sequence Alignment (MSA)^20,21,28,29^, while others do not^30^. Although the set of evolutionary proteins don’t typically have associated measurements for the property of interest, the homology search itself is assumed to implicitly provide a weak, positive labeling of all proteins in the set (*e. g.*, each protein is assumed to have some amount of the desired activity). Thus, we refer to estimating fitness from purely evolutionary data as *weak-positive-only* learning, rather than unsupervised learning.

The second main strategy for estimating ML-based protein fitness models employs supervised learning using protein variants that have had their properties assayed in the laboratory. Depending on the protein and the property of interest, a data set may comprise only hundreds of examples^31^, to hundreds of thousands^32^. The assay may be a relatively direct measurement of the fitness of interest^33^, or be a crude proxy^34^. Additionally, the set of variants may be restricted to one or two mutations away from a query sequence (*e. g.*, a wild type GFP)—a so-called mutational scanning experiment^31,32,34^, or they may be more heterogeneously distributed^33^; the former setting is far more common. The labelled proteins may vary at only a few sequence positions (*e. g.*, four positions^35^), or vary more broadly^32^. By their limited construction, these data sets are not typically believed to be sufficiently rich to comprehensively characterize protein fitness landscapes, but they are a good starting point. As time progresses, the field will train and test on better-powered data sets, revealing increasingly more nuances of fitness landscapes. Nevertheless, increasingly richer supervised models are being applied for the supervised approach, from linear models to convolutional neural networks (CNNs)^12,36^, LSTMs^22,30^, and Transformers^22,23^. Some of these classes of models, such as LSTMs and Transformers, can be employed either as density models or as supervised models. In some cases, a large and broad set of unlabelled proteins is also used to help find useful representations for the supervised model^22,23,26,30^.

There have been recent attempts to combine the two aforementioned approaches, that is, combining the weakly-positive evolutionary data with assay-labelled data^24,26,27^. We refer to such a hybrid learning setting as *weak-positive semi-supervised* learning. Importantly, for many scenarios of practical importance, property measurements can only be made for hundreds of proteins, a regime sometimes referred to as “low-N” protein engineering^26^. Especially in such a regime, but not only, it is important to combine as much useful data as possible so as to learn the most useful protein fitness models. Next we describe existing work that tackles this hybrid problem.

Barrat-Charlaix *et al*.^27^ introduce an elegant approach called an *integrated Potts model* that assumes the energy function of the Potts model for the evolutionary data is identical to that in an energy-based supervised regression model. The two models, coupled by their energy function, are learned jointly. Shamsi *et al*.^24^ instead “transfer” information learned from a first step of modelling evolutionary data to the supervised problem by learning a binary masking of an already-trained evolutionary Potts model using a supervised objective function. Biswas *et al*.^26^ base their approach on the field of Natural Language Processing (NLP)^37,38^, training an unsupervised autoregressive multiplicative LSTM (mLSTM) first on a large database of naturally occurring protein sequences, then refining it with evolutionary data to produce a latent representation called *eUniRep*. This representation is then used as as input features for a regularized linear regression on the assay-labelled data. As such, this eUniRep regression approach uses evolutionary data for the purpose of representation learning, and does not explicitly make use of the sequence density, in contrast to the weak-positive-only learning evolutionary approaches. In general, we call all of these methods *hybrid* approaches because they include both evolutionary and assay-labelled data.

Next we perform a systematic assessment of such hybrid methods, as well as comparisons to methods that use only one of the two data types, and at times also a large reference database of proteins such as is used by *UniRep*. In addition to the methods already developed for combining these data, we also introduce a novel, simple baseline approach, which performs exceptionally well despite its simplicity. In fact, one instantiation of this simple baseline consistently achieves maximal performance on 15 of the 19 data sets used in our assessment—more than any other method—and is typically among the top competitors when it does not. Furthermore, this baseline is a meta-method in that it can be used to augment any purely evolutionary modelling approach to also account for assay-labelled data, and adds low computational burden.

## Results

### Published machine learning methods assessed

We assess a total of 11 machine learning strategies which use evolutionary data, assay-labelled data, or both. Some^23,26,30^ additionally make use of a large “universal” set of unlabelled protein sequences, namely the UniRef50 database^39^. In addition to the three hybrid methods summarized earlier, we also include methods that use only one or the other type of data so as to glean their relative importances across a range of settings. In particular, we include two representative purely evolutionary methods: EVmutation^20^ and DeepSequence^21^, which model the evolutionary sequence density with Potts models and VAEs, respectively. We also include two supervised methods that do not use evolutionary data: a linear regression model with one-hot amino acid features that uses assay-labelled data only, and the ESM-1b Transformer^23^ model which is pre-trained on UniRef50, and then fine-tuned with the assay-labelled data. While several Transformers have been developed and published for protein sequences^22,40^, we chose the ESM-1b model because it has superior performance on a number of prediction tasks when compared to other Transformers^23^. Figure 1 shows a graphical overview of the methods and how they are related.

**Figure 1:**
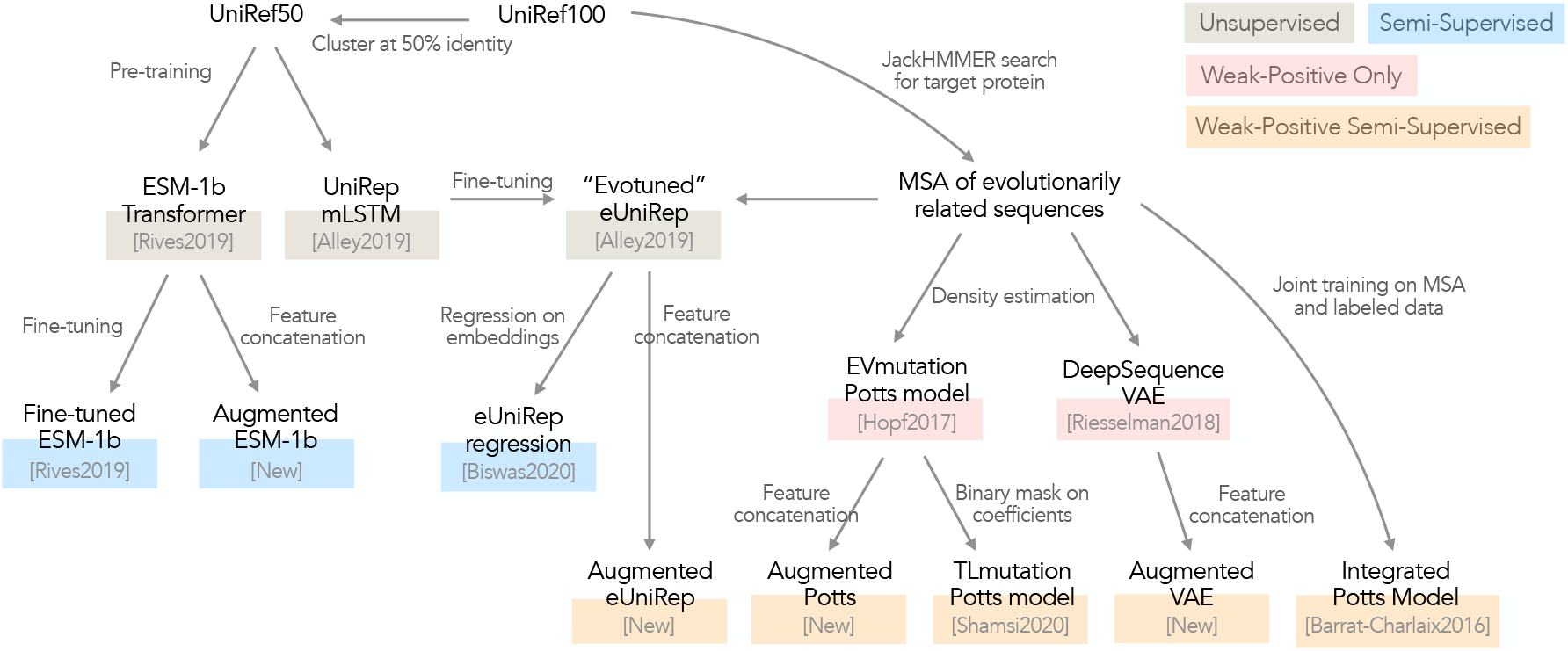
Overview of machine learning strategies evaluated in this paper. We include *weak-positive-only* evolutionary learning approaches (EVmutation^20^ and DeepSequence^21^), semi-supervised learning methods (fine-tuned ESM-1b^23^ and eUniRep regression^26^), and *weak-positive semi-supervised* learning approaches (integrated Potts model^27^, TLmutation^24^, and augmented models).

To enable all of these methods to run on our entire suite of benchmarking data sets we made minor modifications to some of the training procedures so that they could run in a reasonable amount of time. For each modification used, we show that the resulting performance closely matches the performance of the original procedure on data sets used in the original papers (see Methods and Supplementary Figures 3-5).

### A new, simple baseline hybrid method

We developed a new baseline hybrid approach that turns out to be extremely competitive: for any already-trained evolutionary density model, use a linear ridge regression model on one-hot amino acid features augmented with one other feature, namely the sequence density evaluation (Supplementary Figure 1). We refer to this general meta-approach as “augmented”. For example, if the evolutionary density model is a Potts model, we apply our strategy to obtain an *augmented* Potts model. The augmented Potts model can be interpreted in a Bayesian light as updating a prior density model from the evolutionary data with assay-labelled data (see Methods). Because we use only single amino acid features in the regression, effectively we update mainly the single site parameters. If we had included also pairwise amino acid features, then we would be more directly effectively updating all parameters in the Potts model prior, which would yield an approach that is conceptually similar to the integrated Potts model^27^, albeit with different optimization objectives. Our augmenting approach can be readily deployed on any density model, and enjoys a trivial training procedure and correspondingly low computational burden. Although effective ways to incorporate richer features such as pairwise features into the augmented regression could yield further improvements still, we wanted a carefree baseline that did not require feature selection or specialized regularization^41^.

### Assay-labelled data sets

We assess all methods on 19 labelled mutagenesis datasets, each comprising hundreds to tens of thousands of mutant sequences. Most related works^21,23,24,27^ evaluate on a subset of or all of the mutation effect datasets introduced by EVmutation^20^. We included all EVmutation^20^ protein data sets with at least 100 entries in order to have sufficient data to glean insights from. Additionally, we also included a GFP brightness data set^33^ as done by Biswas *et al*.^26^ (their exact data was not available at the time of our assessment). We did not use two other potentially relevant data sets centered on the IgG-binding domain of protein G (GB1)^32,35^ because of the paucity of MSA data available for this protein. In each data set, the mutations are spread across the whole protein sequence or across a particular protein domain. Although most (16 out of the 19) labelled data sets only consist of single mutants that are one mutation away from a fixed wild-type protein, we also include detailed case studies of the three data sets^33,34,42^ with high-order mutants. Supplementary Table 1 provides a detailed overview of these data sets.

### Evolutionary data sets

Each of these labelled data sets is paired with an evolutionary data set found by searching through the UniRef100 database^39^ with the wild-type protein sequence as query. A jackhmmer^43^ search is commonly used for retrieving evolutionary data^20,21,24,26^; however, these search results can differ depending on the jackhammer parameters used. Thus, for simplicity and ease of comparison with other papers^20,21,24^, we use the MSAs provided by EVmutation^20^ when available, and in other cases conduct our own jackhmmer with parameters as used in the EVmutation paper (see Methods).

### Experimental overview

We systematically varied supervised training data sizes by randomly sampling each of 48, 72,···, 216, 240 training examples from each data set. When computationally feasible, we use 5-fold cross-validation for hyper-parameter selection, and otherwise use 20% of the training data. Regardless of the training data size, we always use 20% of the full data (not used for training or validation) as test set. In addition to these fixed sample sizes, for more direct comparison with TLmutation^24^ and ESM^23^, we also test performance using train-test splits, training on 80% of all single mutants in each data set, and testing on the remaining 20% of the data set. For each sample size and the 80-20 split, we evaluate performance over 20 random seeds. We focus on regression fitness models that predict real-valued fitness measures, as opposed to classification models that predict whether a protein sequence is functional or non-functional.

We use two measures of performance: (i) a Spearman correlation between the true and predicted fitness values, and (ii) a ranking measure from the information retrieval community called normalized discounted cumulative gain (NDCG)^12,44^, which gives high (good) values when the top predicted results are enriched for truly high fitness proteins. In reality, the precise metric of relevance is task-specific. For example, in protein engineering, we may be interested only in the best performance among the top *k* proteins that will be assessed in the laboratory. By using both Spearman correlation and NDCG, we hope to be broadly informative across a wide variety of settings. When space does not permit us to include both, the other is always found in the Supplementary Information. Generally, the two metrics yielded similar conclusions.

Results for TLmutation^24^ are omitted from the main figures because the method is on the one hand conceptually similar to our augmented Potts model, but only allows for zeroing out of the evolutionary Potts model parameters by way of supervised learning, and because it performed worse than our augmented Potts model (see Supplementary Figure 2). Moreover, TLmutation was much more computationally expensive to run than the augmented Potts model.

### Comparison of existing methods and the augmented Potts model

When comparing existing methods to each other and to our augmented Potts model, averaged over all data sets, the augmented Potts model typically outperformed all other methods, including more sophisticated methods such as eUniRep regression^26^ (Figure 2). The sole exception was that the purely evolutionarily-based VAE outperformed the augmented Potts model on the double mutant data in the low data regime. However, later we will show that augmenting the VAE itself improves the VAE results further, suggesting that the Potts model is not sufficiently rich in some cases. As expected, with increasing labelled training data, the hybrid and purely supervised methods increase in performance, whereas the purely evolutionary approaches remains constant. When breaking down the average performance into individual data sets, the augmented Potts model is competitive on a majority of data sets (Supplementary Figures 10-13). Remarkably, the extremely simple supervised-only method of linear regression on one-hot amino acid features is also among the top performers in the 80-20 split setting.

**Figure 2:**
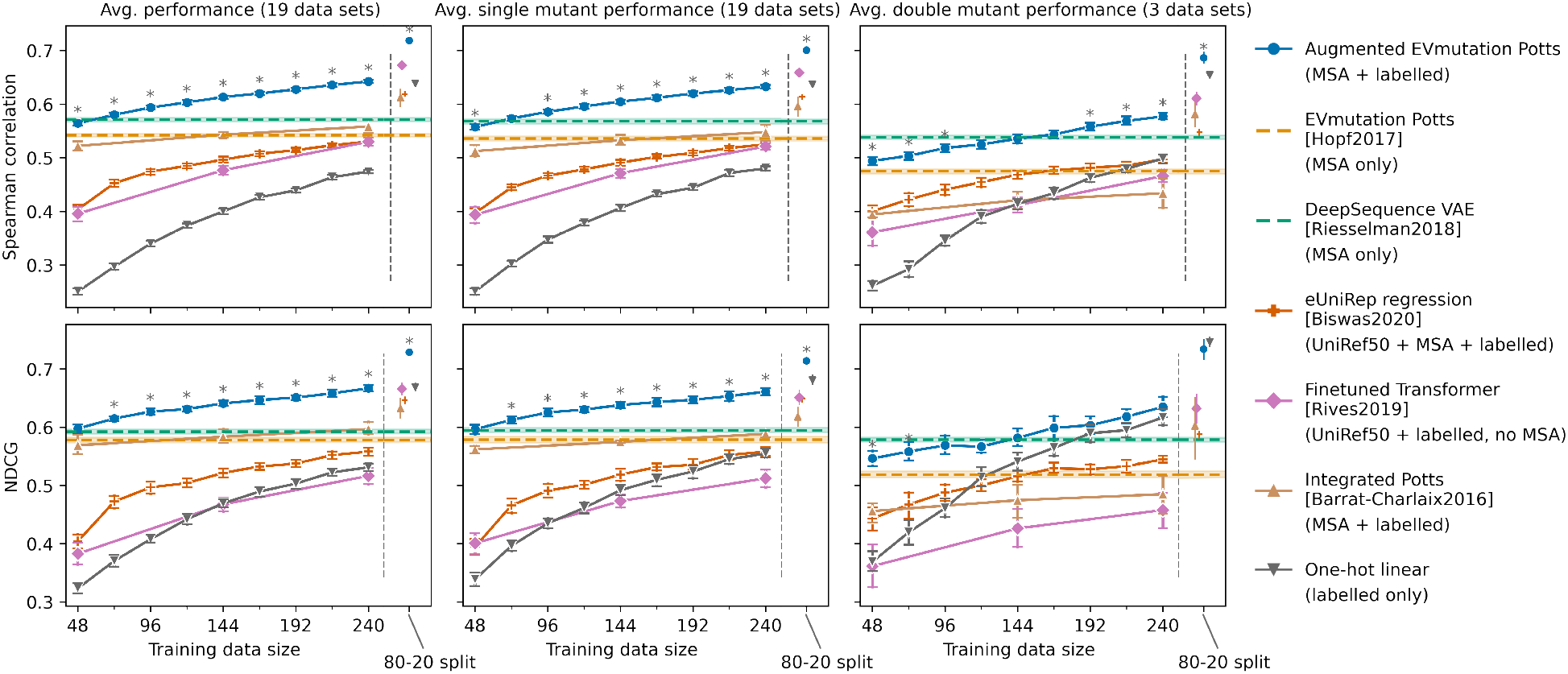
Comparison of existing machine learning approaches and the augmented Potts model. Spearman correlations and normalized average discounted cumulative gains (NDCG) are averaged over data sets, with error bars indicate bootstrapped 95% confidence intervals estimated from 20 random seeds for sampling training and test data. The horizontal axis shows the number of supervised training examples used, or else when the 80-20 train-test split was used. The left column indicates performance on the full test data, while the middle and right columns indicate performance among all single and double mutants in the test data. Asterisks (*) indicate statistical significance (*p* < 0.01) from a two-sided Mann-Whitney U test that the augmented Potts model is better than every other method. See Supplementary Figure 8-13 for breakdown.

Due to limited availability of data sets that include double mutants, the reported double mutant performance is averaged over only the three relevant data sets and hence is heavily influenced by their characteristics. We refer the reader to Supplementary Figure 8-9 for the double mutant performance breakdown.

### Augmented models as a general, simple, and effective strategy

We next tried augmenting not only the EVmutation Potts model, but also the DeepSequence VAE, eUniRep mLSTM, and ESM-1b Transformer. Overall, we found that the augmented VAE achieved the highest average performance among the augmented models, with the augmented Potts model a close second (Figure 3). No matter which density model we augmented, the augmented version of the model always improved the performance, regardless of the training data set size.

**Figure 3:**
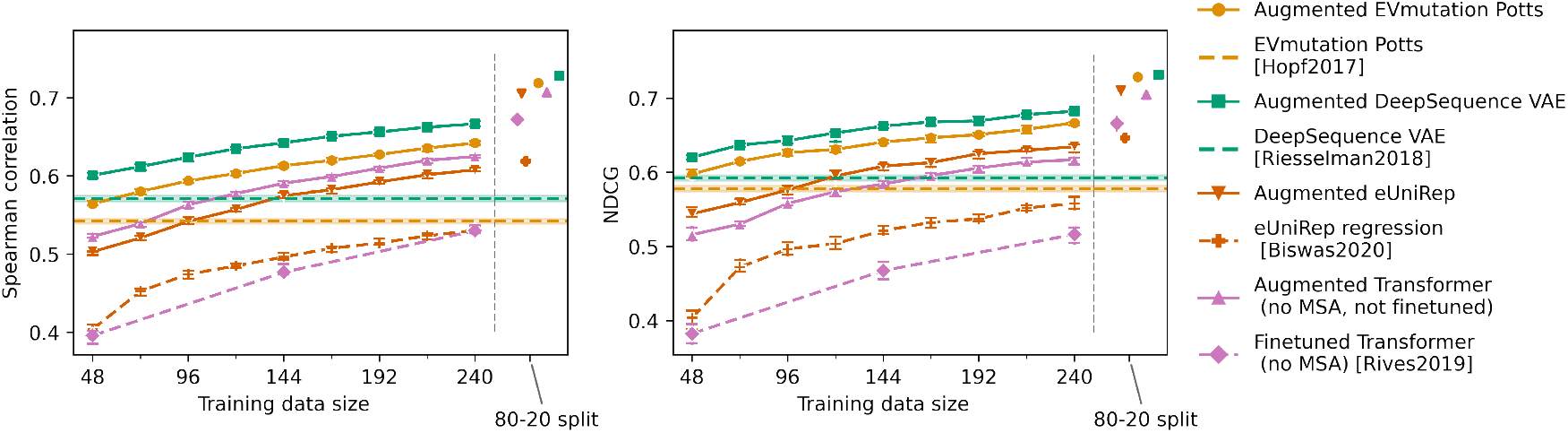
Comparison of augmented density models. Dashed lines show baseline evolutionary density model performance, while solid lines in the matching color show the corresponding augmented model. The evaluation setup here is identical to the left column in Figure 2.

Next we counted how many times any given approach, among all approaches evaluated so far, tended to be the best performer (Figure 4). As soon as any labelled data are made available in addition to evolutionary data, the augmented VAE wins the most, followed by the other augmented models which are roughly comparable—other than the augmented eUniRep which underperforms with only 48 training data points. In all the experiments so far VAEs tend to perform slightly better than Potts models (when both are augmented, and when neither is augmented) on average. However, it is worth noting that the VAE does do worse on the GAL4 DNA-binding domain and on the kanamycin kinase APH(3’)II. This observation is consistent with the comparison between VAEs and Potts models in Riesselman *et al*.^21^’s Figure 3.

**Figure 4:**
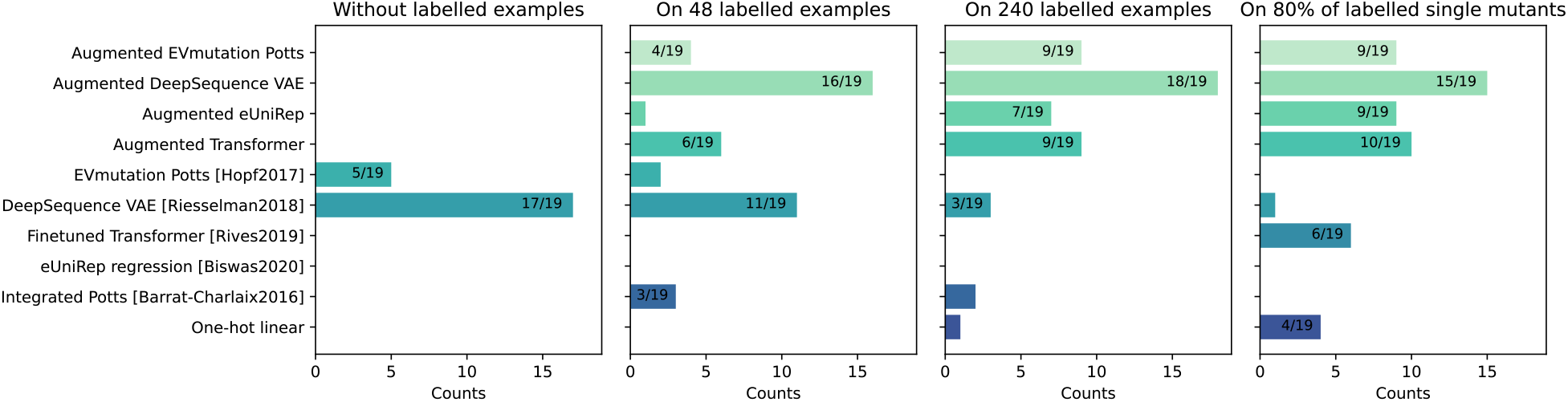
Frequency of each method achieving the highest Spearman correlation per data set. The horizontal axis shows the number of times a given modelling strategy had better Spearman correlation than all other methods. This count is computed by first identifying the top-performing method for any given scenario, and then also including any other methods that come within the 95% confidence interval of this top performer. Then both the top performer and those within the 95% confidence interval are counted. Four settings are used: with no labelled data, when training on 48 or 240 labelled single-mutant examples, and in the 80-20 train-test split setting. See Supplementary Figure 14 for the normalized discounted cumulative gains (NDCG) version of this figure.

### Extrapolation from single to higher-order mutants

Because 16 of the 19 supervised data sets consisted of only single mutants of a wild type, we next more closely examined the three data sets with higher-order mutants, namely, data on the green fluorescent protein (GFP)^33^, the Poly(A)-binding protein (PABP) RRM domain^34^, and the ubiquitination factor E4B (UBE4B) U-box domain^42^. In each case, we trained the models using only single-mutant labelled data, and then evaluated on all the other mutants (including single, double, triple, and quadruple mutants, as shown in Supplementary Figure 8). The results here echo the results of our earlier comparisons (Figure 2 and Figure 3) in the sense that the augmented Potts model and the augmented VAE outperform existing methods. It is worth noting, however, that on the E3 ligase ubiquitination factor E4B (UBE4B), none of the methods achieve good performance on higher order mutants. Such poor performance could be due to poor correlation between fitness in evolution and the selective pressure in the assay. If indeed this is the case, then hybrid methods should be able to effectively discount evolutionary data as increasingly more supervised data becomes available. While it is often difficult to know *a priori* whether evolutionary fitness correlates well with the property-of-interest, if one knew that the correlation was not useful, one would clearly use a purely supervised approach.

In data sets where the assay-labelled variants comprise a variety of edit distances from the wild type, it is interesting to observe that simply using the edit distance itself can be highly predictive of fitness. For example, for GFP fluorescence, edit distance as a predictive model achieves a Spearman correlation of 0.447 (Figure 5), presumably because most mutations are deleterious and roughly additively so. This result suggests that correlation measures can be driven by rather simple features, and that by explicitly using these in simple settings, one can better deduce from where the predictive signal is arising. In contrast to GFP, mutation count is not as strongly predictive for the UBE4B U-box domain data (Supplementary Figure 15), presumably because the mutational effects are more heterogeneous in the sense that they comprise either more diverse (both deleterious and beneficial), or non-additive effects, or both. Moreover, none of the other methods perform particularly well on the UBE4B U-box domain data (Supplementary Figure 8-9 and 15). Indeed, the more heterogenous the mutational effects, the more rugged the fitness landscape, requiring increasingly more diverse and larger training data to model well.

**Figure 5:**
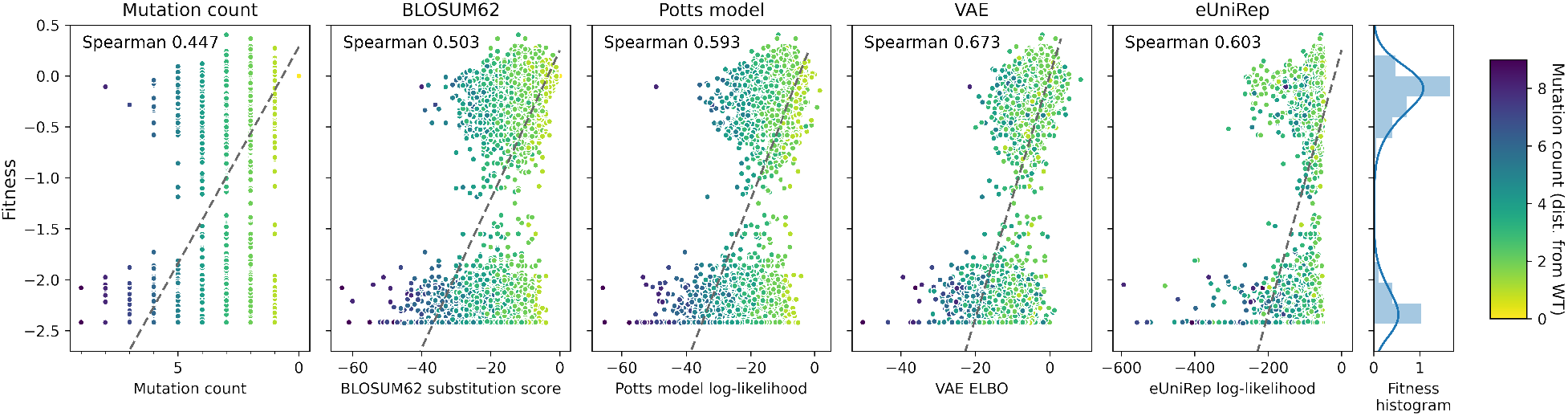
Mutational count as a predictive model on GFP. Each dot represents a GFP sequence, with darker color indicating further distance from the wild-type. Even the naive baseline of a single integer mutation count, is highly rank-correlated with experimentally measured brightness. Supplementary Figure 15 shows a similar plot for the UBE4B U-box domain.

## Discussion

We presented the first systematic comparison of hybrid machine learning strategies for protein fitness prediction. That is, we investigated methods that make use of both evolutionary and assay-labelled data. In addition to comparing to published approaches, we introduced a new baseline, the *augmented* density model meta-approach, which we instantiated here through the augmented Potts model, augmented VAE, augmented eUniRep, and augmented Transformer. Across a wide range of settings, one of the simplest possible models—the augmented Potts model—is competitive with far more complicated and computationally expensive deep-learning-based approaches, such as Transformer- and LSTM-based approaches. Indeed, in the relatively larger data settings—which were still quite small—even a simple linear regression using one-hot amino acid encoding performed quite well. This baseline has sometimes been neglected in comparisons of vastly more complicated methods. Overall, the augmented VAE performs best. When thousands of labelled examples are available, the augmented ESM-1b Transformer was competitive with other methods despite not relying on evolutionary data at all, making it a promising choice for proteins with limited evolutionary data and substantial labelled training data. Furthermore, faster and more accurate approximations to the Transformer sequence log-likelihood have the potential to improve performance further. The fact that the augmented Transformer outperforms the fine-tuned Transformer suggests it may be worth revisiting some of the conventional embedding and fine-tuning techniques from NLP when using them for tasks that include weak-positively labelled data.

As larger and more variable assay-labelled data sets emerge, we expect that increasingly richer models will dominate. Moreover, different machine learning strategies are likely needed for the “low-N”, or limited labelled data, regime and for the “high-N” regime, both of which are likely to remain. Therefore, when making general methodological recommendations, researchers should either consider implications across the full range of labelled data sizes or restrict recommendations to the regime tested.

Because the comparisons herein were already vast, we did not attempt to bring auxiliary information into our evaluation. Recent studies^12,13,45–47^ show that protein structures, Rosetta energy terms, and molecular dynamics also provide valuable information in addition to protein sequence information. It therefore stands to reason that such features are likely to be helpful when included in any approach that allows for them. However, we anticipate that the overall conclusions of our comparison are likely to remain in such a context. Importantly, as more features go into the augmentation, there are likely opportunities to use a more complex modelling strategy in the augmentation, based on regression trees, deep learning, or other feature-construction approaches.

Although our goal was to assess different methods on real data rather than assuming certain simulation settings are representative, these real data sets have their limitations and are not necessarily representative of the test sets that one might encounter for protein engineering. In particular, in a protein engineering setting, one would like to understand how the model performs across a larger swath of protein space. Most importantly, one would like predictive models to extrapolate as accurately as possible to higher fitness areas than that observed in the data^48^, and to be able to gauge when such extrapolations are unreliable^17^. While it is possible to attempt to tackle extrapolation using techniques from covariate shift such as importance-weighted empirical risk minimization^48,49^, such approaches come with their own problems, and hence we focused on directly evaluating methods without such corrections to account for domain shifts.

Another issue we did not explore is to what extent a particular protein’s fitness is correlated with the density of a corresponding set of evolutionary sequences. Herein, we focused predominantly on proteins for which some such correlation has already been shown to exist, and further assumed that the extent of this correlation did not differentially affect the methods assessed.

Finally, while we have focused on evaluating predictive performance, in cases where the predictive model is used in conjunction with a design strategy^12,17–19,48^, it could be useful to evaluate both the outcome of the design process, and the predictive model by itself, rather than tying them up into a single comparison, so as to achieve a finer-grained understanding.

## Methods

### Evolutionary sequences

We use the MSAs provided by EVmutation^1^ whenever possible. For the green fluorescent protein (GFP), the only exception, we follow the same procedure as EVmutation to gather sequences using the profile HMM homology search tool jackhmmer^2^. We determine the bit score threshold in jackhmmer search with the same criterion from EVmutation. In particular, for GFP, we started with 0.5 bits/residue and subsequently lowered the threshold to 0.1 bits/residue to meet the sequence number requirement (redundancy-reduced number of sequences ≥ 10*L* where *L* is the length of the aligned region). For sensitivity analysis, when using the bit score to 0.5 bits/residue or increasing the number of iterations to 10, the resulting MSAs on GFP still lead to similar downstream model performance (data not shown).

### Mutation effect data sets

Hopf *et al*.^1^ identified a list of mutation effect data sets generated by mutagenesis experiments of entire proteins, protein domains, and RNA molecules. We exclude the data sets for RNA molecules and influenza virus sequences, as well as excluding data sets that contain fewer than 100 entries, in order to have meaningful train/test splits with at least 20 examples in test data. This leaves us with 18 data sets from EVmutation.

Following the convention in EVmutation and DeepSequence, we exclude sequences with mutations at positions that have more than 30% gaps in MSAs, to focus on regions with sufficient evolutionary data. On most data sets, this excludes less than 10% of the data, although for a few proteins such as GFP this affects as much as half of the positions. For example, on GFP, out of 237 positions, only positions 15-150 pass the criterion of less than 30% gaps in the MSA. Coincidentally, the selected position 15-150 region covers the 81 amino region studied by Biswas *et al*.^3^.

Among those 18 EVmutation data sets, only two on the poly(A)-binding protein activity^4^ and the UBE4B auto-ubiquitination activity^5^ include higher-order mutants.^†^ While EVmutation only evaluates performance on the single mutants from the UBE4B U-box domain data^5^, we also include higher-order mutants in evaluation. Additionally, we also include a green fluorescent protein (GFP) brightness data set^7^, since GFP is a core example used by Biswas *et al*.^3^. The GFP data also include higher-order mutants. This results in 19 data sets total as listed in Supplementary Table 1.

### Encodings

In addition to the one-hot amino acid encoding, we also experiment with the 19-dimensional physicochemical representation of the amino acid space developed by Georgiev^8^, as also used by Wittman *et al*.^9^. The features in Georgiev representation are principal components of over 500 amino acid indices from the AAIndex database^10^. As shown in Supplementary Figure 7, the Georgiev encoding achieves better performance than the one-hot amino acid encoding on data sets with higher-order mutants when presented with sufficient training data (bottom right), agreeing with previous findings^9^. However, on single-mutant only data sets or on limited training examples, the Georgiev encoding leads to almost identical performance as the one-hot encoding. For the augmented Potts model, the Georgiev encoding also does not improve performance compared to the one-hot encoding in any setting.

Since the other evaluated methods, such as EVmutation^1^, DeepSequence^11^, and UniRep^12^, all use the one-hot amino acid encoding as inputs to their models, we omit the Georgiev encoding results in the main text for fair comparison.

### One-hot linear model

For a sequence *s* = (*s*_1_,···, *s*_*L*_) of length *L* in alphabet 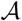 where 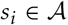, the linear model with one-hot amino acid encoding maps the sequence to a scalar regression output by

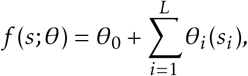

where *θ*_0_ ∈ ℝ is the bias term, and each 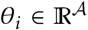 is a 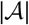-dimensional vector. Each coefficient of *θ*_i_ corresponds to a different amino acid at position *i*, and we use the notation *θ*_i_(*s*_i_) to denote the coefficient corresponding to the particular amino acid *s*_i_. When fitting the linear model, we use ridge regression with regularization strength chosen by five-fold cross validation.

Importantly, when training and testing the one-hot linear model on non-overlapping single mutants, only the coefficients corresponding to wild-type amino acids appear in both training and testing, and therefore effectively the model can only learn position-specific information from mutations but not position-specific amino-acid specific information on single-mutant-only data sets.

### EVmutation Potts model

Briefly, EVmutation learns a sequence distribution under site-specific constraints and pairwise constraints for each protein family. Under a Potts model (also sometimes known as a generalized Ising model) with alphabet 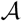, the probability of a sequence 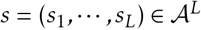 of length *L* is given by a Boltzmann distribution:

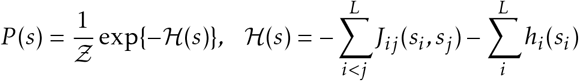

where *Ƶ* is a normalization constant, 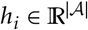 are site-specific amino-acid-specific parameters, 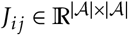 are pair-specific amino-acid-specific parameters, and 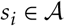 indicates the amino acid at position *i*. The *h* and *J* parameters are estimated by regularized maximum pseudo-likelihood. See EVmutation paper for the full modeling fitting details of sequence re-weighting, pseudolikelihood approximation, regularization, and optimization. We use the same plmc package as EVmutation to fit Potts models with default parameters.

Under the assumption that learned sequence probabilities are correlated with sequence fitness, EVmutation predicts the mutation effect of a mutant according to the log-odds ratios of sequence probabilities between the wild-type and mutant sequences, which is equivalent to the log-likelihood of the mutant up to a constant.

### DeepSequence VAE

Similar to EVmutation, DeepSequence^11^ also models a sequence distribution for each protein family and predicts mutation effects according to approximations of log-odds ratios of sequence probabilities between the wild-type and mutant sequences. Here, the sequence distribution is modeled by a nonlinear variational autoencoder model with a multivariate Gaussian latent variable *z*. A neural network parameterizes the conditional distribution *p*(*s* | *z, θ*), and the sequence likelihood *p*(*s* | *θ*) is then

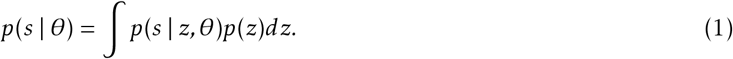

While the exact log likelihoods are intractable, they are lower-bounded by the evidence lower bound (ELBO)

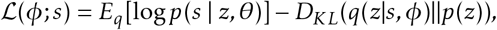

where *q*(*z* | *s,ϕ*) is a variational approximation for the posterior distribution *p*(*z* | *s, θ*) that is also modeled by a neural network. As an approximation to log-likelihoods, the ELBO can also be used to predict mutation effects. Whenever possible, we use the available ensemble predictions from DeepSequence^11^. For the GFP, the UBE4B U-box domain, and the BRCA1 ring domain, we followed the DeepSequence code and parameters to train an ensemble of VAE models (5 models with different random seeds). When computing the evidence-based lower bound (ELBO) as sequence log-likelihood estimations, we follow DeepSequence to take the average of 2000 ELBO samples (400 samples from each of the 5 VAE models in the ensemble).

### eUniRep mLSTM

The UniRep^12^ multiplicative long short-term memory network (mLSTM) can also be viewed as a sequence distribution. Given the first *i* amino acids of a sequence, a neural network parameterizes the conditional probabilities of the next amino acid *P*(*s*_*i*+1_ | *s*_1_ ··· *s*_i_). From the conditional probabilities we can also reconstruct the sequence probability as

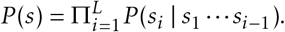

For convenience, artificial start and stop tokens are added at the two ends of sequences. Under this probabilistic interpretation, the unsupervised pre-training on UniRef50 sequences with next-step predictions is equivalent to finding model parameters that maximize likelihood on UniRef50 sequences. We omit the performance of the pre-trained (not evo-tuned) UniRep model in our results since generally it is largely outperformed by the eUniRep described below.

To adapt the UniRep model to specific protein families, Biswas *et al*.^3^ propose the “evo-tuning” procedure, which fine-tunes the UniRep model by minimizing the same next-step prediction loss on evolutionary sequences in addition to on UniRef50 sequences. Following the naming convention^3^, We refer to the model after “evo-tuning” as eUniRep. After fine-tuning, Biswas *et al*.^3^ then perform supervised regression with the 1900-dimensional average embeddings (averaged over sequence length axis) from the final hidden layer in the eUniRep model as regression features.

While the original fine-tuning procedure for eUniRep^3^ optimizes for next-step prediction loss on entire evolutionary sequences up to length 500 (discarding longer sequences), we find that next-step prediction on aligned portions of evolutionary sequences (with gaps included as gap tokens) works as well on GFP and beta-lactamase (Supplementary Figure 4), and therefore fine-tune on only aligned portions to lower computational costs as the mLSTM memory usage scales quadratically with respect to sequence length.

For each protein family, we randomly split the evolutionary sequences from the MSA into an 80% training set and a 20% validation set to check for over-fitting. When training for 10,000 gradient steps with the same learning rate 1 × 10^−5^ as Biswas *et al*.^3^, the validation loss eventually plateaus but does not increase. Since the validation loss typically plateaus before 10,000 steps, we chose to stop training at 10,000 steps.

### ESM-1b Transformer

The ESM-1b Transformer^13^ is pre-trained on UniRef50 representative sequences with the masked language modeling objective, where a fraction (15%) of amino acids in each input sequence are masked and the model is trained to predict the missing tokens. We chose the ESM-1b model to represent Transformers as it has the best performance on downstream secondary structure prediction and contact prediction tasks^13^ among Transformers from prior works^14,15^ and other Transformer architectures tested. While the original mutation effect prediction results^13^ were based on the older 34-layer ESM-1 model, we found that ESM-1b slightly improves performance over the ESM-1 model (data not shown).

When using Transformer models for sequence fitness predictions, Rives *et al*.^13^ mask the mutated positions and used the difference in conditional log-likelihoods (conditioned on non-mutated amino acids) between the mutated amino acids and the wild-type amino acids as fitness prediction. We explain below that this could be viewed as an approximation for pseudo log-likelihoods (PLLs).

Masked token language models can also be viewed as sequence distributions, where the sequence distribution is implicitly represented by conditional likelihoods *P*(*s_i_* | *s*_−*i*_), where *s*_−*i*_ indicates all other sequence positions excluding position *i*, i.e. *s*_−*i*_ = *s*_1_ ··· *s*_*i*−1_*s*_*i*+1_ ··· *s_L_*.

Since exact likelihood are too computationally expensive for Transformers, we resort to pseudo-likelihoods. In general, given a sequence *s* of length *L*, its pseudo-likelihood^16^ is defined as the product of conditional likelihoods for each site. Hence, the pseudo-log-likelihood of a sequence *s* is

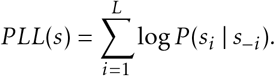

However, even the evaluation of pseudo log-likelihoods (PLLs) is computationally expensive, since it requires *L* inferences for a sequence of length *L*. As a more computationally efficient approximation, we only compute conditional likelihoods on the mutated positions, and then use the difference between the conditional log-likelihoods of the mutated sequence and the wild-type sequence as an approximation for pseudo-log-likelihoods. More specifically, for a mutant sequence *ϕ* and wild-type sequence *σ*, we approximate the PLL difference by the following, as equivalent to the existing formula used by Rives *et al*.^13^:

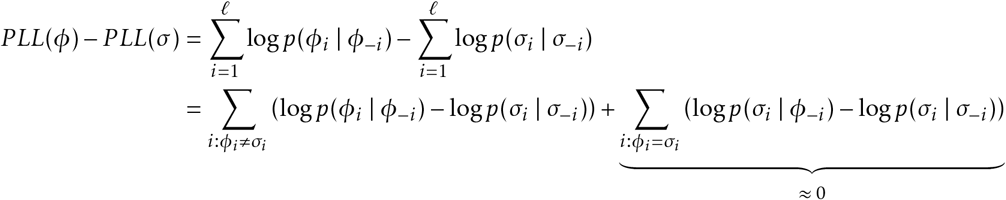

The underlying assumption here is that the conditional log-likelihoods of wild-type amino acids on mutant backgrounds are roughly the same as on wild-type backgrounds. While this might be accurate for mutant sequences that are close enough to wild-type sequences, in general there is no support for this approximation on high-order mutant sequences and the full pseudo-log-likelihoods will likely be more accurate for mutation effect predictions.

For supervised learning, we evaluated two approaches with the same Transformer model. The first one (“fine-tuned Transformer”) is to fine-tune the entire Transformer model to regress the PLL difference to mutation effects as done by Rives *et al*.^13^. We perform 20 epochs of supervised fine-tuning with learning rate 3 × 10^−5^ and with early stopping according to validation Spearman correlation. For a given training data size, we keep 20% of the data for validation (early-stopping) and use the remaining 80% for fine-tuning. Using Spearman correlation as opposed to validation loss for early stopping is crucial for performance, especially on small data sizes. Although the implementation details might not exactly match those from Rives *et al*. (since the fine-tuning code is not available), we show that our method is able to reproduce the same results on most of the Envision data sets used by Rives *et al*., as shown in Supplementary Figure 3.

In the second approach (“augmented Transformer”), while keeping the Transformer model constant, we concatenate the PLL difference inferred from the pre-trained (not fine-tuned) model together with one-hot amino acid encoding as features for regression.

We also attempted unsupervised fine-tuning (“evo-tuning”) of the ESM-1b Transformer model on evolutionarily related sequences from MSAs, although our preliminary efforts on *β*-lactamase and PABP-RRM do not result in improved performance (data not shown).

### Integrated (tied-energy) Potts model

The integrative approach^17^ for Potts models optimizes the joint log-likelihood for model parameters on both evolutionary data and labelled data. Given evolutionary sequences *σ*_1_,···, *σ_M_*, the log-likelihood on evolutionary data is

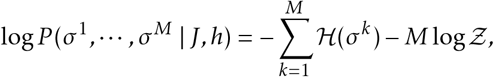

where

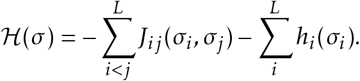

On the other hand, given sequence-fitness pairs (*s*_1_, *y*_1_),···, (*s*_*N*_, *y_N_*), assuming i.i.d. Gaussian noise drawn from 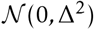, the log-likelihood on labelled data is

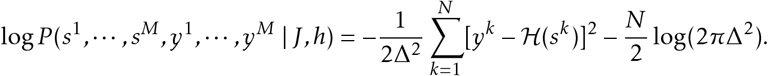

The integrated Potts model is learned by maximizing the joint log-likelihood log *P*(*σ*^1^, ···, *σ^M^* | *J, h*)+log *P*(*s*^1^, ···, *s^M^*,*y*^1^, ···,*y^M^ J, h*) with *ℓ*_2_-regularization.

The noise variance Δ^2^ determines the relative weighting between the two losses in the joint log-likelihood. Using existing notation^17^, the relative weighting parameter is

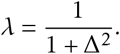

In practice, since we do not know the noise variance, we use 20% of the training data for validation and choose the best *λ* according to Spearman correlation on validation set. Another practical complication is that we only have fitness labels that are up to some monotonic transformation from the energies. Following Figliuzzi *et al*.^18^, we use a monotonic mapping between sorted energy values and sorted fitness labels.

Following recommendations from the authors, the Potts model parameters are initialized with parameters estimated by pseudo-likelihood. Before introducing labelled data, we first give the model a warm start by training 500 iterations only on evolutionary data. Then, we optimize the regularized joint log-likelihood for 100 iteration for each *λ* value. The only exception is for *β*-glucosidase, where the protein sequences are too long (> 500 amino acids) and lead to very high memory consumption and long run-time (> 2 days) for the integrated Potts model. Therefore, for *β*-glucosidase alone we use the Potts model performance without labelled data as a substitute for the integrated model performance.

We follow the publicly available code (https://github.com/PierreBarrat/DCATools/tree/master/src) with slight modifications to use zero-sum gauge instead of wild-type gauge for Potts models. In Potts models for categorical variables, there are more free parameters than independent constraints, and gauge fixing refers to reducing the number of independent parameters to match match the number of independent constraints^19^. The wild-type gauge forces all parameters corresponding to wild-type amino acids to be zero, while the zero-sum gauge requires 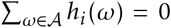 and 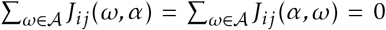. When there is regularization involved, different gauge fixings lead to non-equivalent models. Although gradient calculations are easier in the wild-type gauge, we found that the zero-sum gauge led to better predictive performance (Supplementary Figure 5) and hence adopted the zero-sum gauge.

### Alternative tied-energy models

In addition to integrated Potts models, we also evaluate alternative ways to train a density model of a protein family and a predictive model of fitness in an integrated fashion. Rather than using a fixed and possibly incorrect monotonic mapping between sorted energy values and sorted fitness labels^18^, we consider a differentiable proxy of the Spearman correlation^20^ between the predicted and true fitness values. Specifically, given *N* sequence-fitness pairs, (*s*_1_, *y*_1_), ···, (*s_N_*, *y_N_*), let (*s*_1_,*r*(*y*_1_)),···, (*s_N_*,*r*(*y_N_*)) denote the corresponding sequence-ranking pairs, where *r*(*y_k_*) ∈ {1,…, *N*} gives the rank in descending order of the value *y_k_* among the values {*y*_1_,…, *y_N_*}. We fit a Potts model by optimizing the joint loss

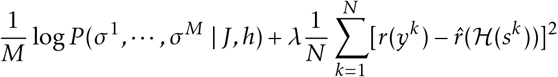

with *ℓ*_2_-regularization, where 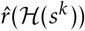 is a differentiable proxy^20^ of the rank 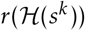. For tractability, we optimize the pseudo-likelihood approximation of the log-likelihood term in the joint loss. We choose the value of *λ* ∈ {0.0001, 0.001, 0.01, 0.1, 0.9} with 5-fold cross-validation.

To explore the effect of training energy-based models (EBMs) other than the Potts model in this fashion, we use the same procedure to fit an EBM with an energy function parameterized by a two-layer feed-forward neural network with hidden layer sizes of (300, 100) and ELU activations. As with the Potts model, we optimize the pseudo-likelihood approximation of the log-likelihood term in the joint loss.

### BLOSUM62 substitution scores

For a mutant sequence, we sum the BLOSUM62 substitution scores between every amino acid in the wild-type sequence and the mutant sequence. The scores are then calibrated up to a constant to be zero for the wild-type sequence.

### Augmented models

To augment an existing density model, we concatenate the sequence density estimation from the original model together with one-hot encoding as features, as illustrated in Supplementary Figure 1. Mathematically, following the same notation for one-hot linear models, given a density model with sequence probability distribution *p*(*s*; *φ*) and fixed density model parameters *φ*, the corresponding augmented model is

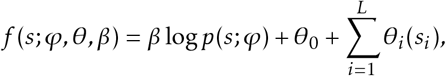

where the parameters 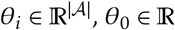, and *β* ∈ ℝ are learned from assay-labelled data. In Ridge regression on the joint features, the regularization strength for the coefficient *β* for sequence density estimation is set to 10^−8^ (practically unregularized, set to a very small quantity for ease of implementation), while the one-hot encoding parameters *θ* are regularized with cross-validated regularization strength. For density models where computing exact log-likelihoods is challenging, we use pseudo-log-likelihood instead of loglikelihood as a regression feature.

### Bayesian interpretation of augmented Potts model

The augmented Potts model can also been seen as updating a Bayesian Potts model prior with assay-labelled data. If we fit the augmented Potts model in a two-step procedure, first fitting a scalar parameter *β* for the log-likelihood feature and then fitting the remaining parameters, then the second ridge regression estimate step effectively fits a one-hot linear model to the residuals *y′* = *y* − *β* log*p*(*s;φ*). Under the Bayesian interpretation of ridge regression, this is equivalent to a zero-mean Gaussian Bayesian prior 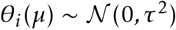 on all one-hot coefficients *θ_i_*(*μ*) (coefficients for regressing on the residuals) for i = 1, ··· , *L* and 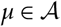, where *τ* is determined by the regularization strength. Therefore, the augmented Potts model can be seen as a Bayesian update on site-specific parameters *h_i_* where the prior

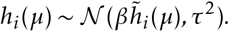

is centered on a scaled version of the site-specific Potts model parameter 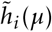 estimated from evolutionary data. Intuitively, the augmented Potts model keeps the pairwise Potts model parameters fixed while correcting for the site-specific parameters according to labelled data. In the most general formulation, one could also further augment Potts models with pairwise features (not shown).

### Model evaluation

For each data set, we randomly sample 20% of the data set as held-out test data. Among the remaining 80% data, we randomly sample *N* = 24, 48, 72, 96, ···, or 240 single mutant sequences as training data in separate experiments, or use all single mutant sequences in the 80% training data in the 80-20 split experiments. When computationally feasible (for one-hot linear model, eUniRep regression, and all augmented models), we use five-fold cross-validation to determine hyperparameters. Otherwise, for the fine-tuned Transformer and the integrated Potts model, we set aside 20% of the training data to determine hyperparameters (*i. e.,* the number of fine-tuning epochs for the Transformer and the relative weighting between evolutionary and assay-labelled data for the integrated Potts model). For each fixed sample size, the model evaluation procedure is repeated 20 times with different random seeds for uncertainty estimation. The only exception is for the integrated Potts model, where we only use 5 random seeds due to the long computation time. The 95% confidence intervals of the means are estimated via bootstrap-ping.

### NDCG

Discounted cumulative gain (DCG) is a common measure of ranking quality. When sorting all examples by predicted score 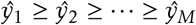, DCG sums the true scores *y* after applying a logarithmic discount,

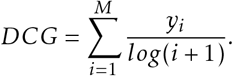

When calculating the DCG, we first standardize the true scores *y* to have zero mean and unit variance to make DCG values comparable across data sets. The normalized discounted cumulative gain (NDCG) metric further normalizes DCG by dividing by the best possible score (*i. e.,* ideal DCG, obtained for a perfect ranking) to obtain a score between 0 and 1. The NDCG score can be seen as a smoothed version of the mean label of the top *k* predictions, without having to fix an arbitrary *k*.

### Mann–Whitney U test

We use the two-sided nonparametric Mann–Whitney U test for comparing average performance between different methods. The null hypothesis of the U-test is that, for randomly selected values *X* and *Y* from two populations, the probability of *X* being greater than *Y* is equal to the probability of *Y* being greater than *X*. The alternative hypothesis is that one population is stochastically greater than the other. In the context of Figure 2, since we are comparing the average performance over all data sets, the population consists of the average performance of a given method computed from different random seeds. This satisfies the assumption that the observations in each population are independent of each other.

## Data availability

All data used were publicly available through citations available in the paper. A processed version of the data used herein will be made available upon publication.

## Code availability

The evaluation framework, method implementations, and analysis code will be made publicly available upon publication.

## Acknowledgments

We thank A. Aghazadeh, P. Almhjell, F. Arnold, A. Busia, D. Brookes, M. Jagota, K. Johnston, L. Schaus, N. Thomas, Y. Wang, and B. Wittmann for helpful discussions. We would also like to thank P. Barrat-Charlaix, S. Biswas, J. Meier, and Z. Shamsi for providing helpful details about their methods and implementations.

## Footnotes

† Even though EVmutation Supplemental Table 1 indicates that the YAP1 (WW domain 1) peptide binding data set also include higher-order mutants, the supplemental data for YAP1 only contain single-mutant sequences, and no higher-order mutant data are available in the original publication6 either.

