## Supplementary information for "Combining evolutionary and assay-labelled data for protein fitness prediction"

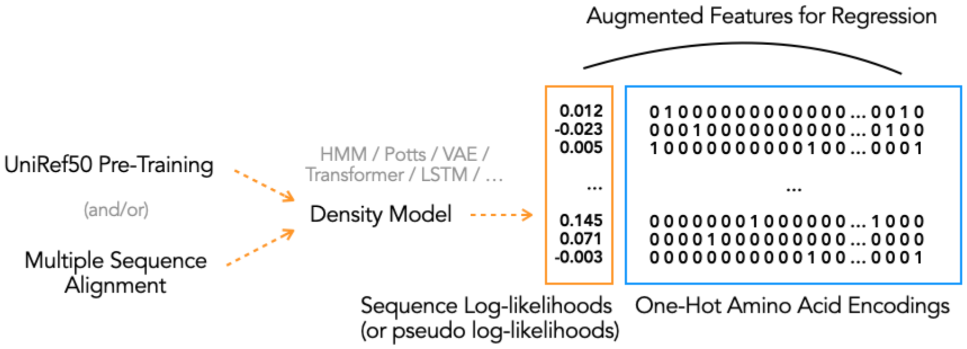


Supplementary Figure 1: **Illustration of augmented models.** In addition to evaluating existing machine learning strategies from the literature, we also propose feature concatenation as a simple meta-approach to combine evolutionary and assay-labelled data. Based on any sequence density model learned from evolutionary sequences, we augment the inferred sequence log-likelihoods or pseudo log-likelihoods with additional one-hot amino acid encoding features for regression.

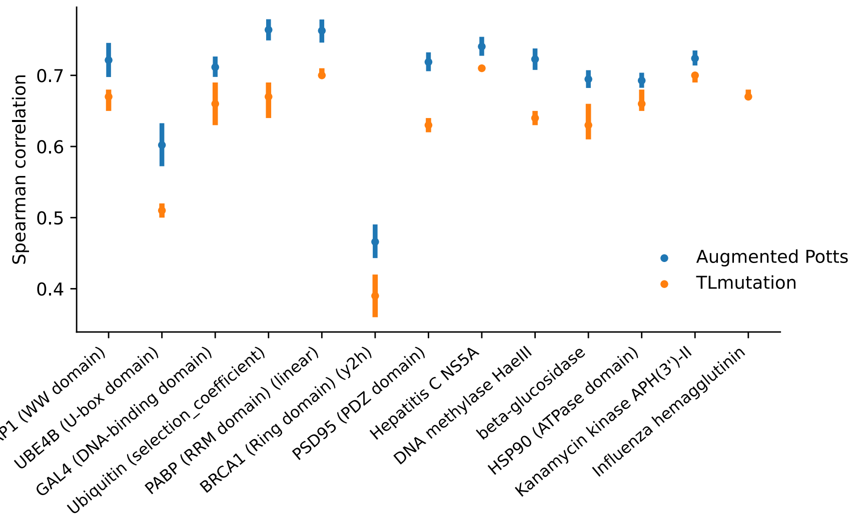


Supplementary Figure 2: **Comparison of TLmutation and the augmented Potts model**. While both TLmutation and augmented Potts models seek to update Potts model parameters with assay-labelled data, augmented Potts models learn additional site-specific parameters to correct the Potts model, whereas TLmutation learns a binary 0/1 mask on the Potts model parameters. When comparing the two methods with random 80-20 train-test splits on the data sets used in TLmutation, the augmented Potts model achieves higher Spearman correlation on most data sets. The TLmutation performance shown is based on Supplementary Figure S1 from Shamsi et al.[1]. The $\beta$-lactamase entry from Supplementary Figure S1[1] is omitted here as it is unclear which $\beta$-lactamase feature column from the data set is referred to.

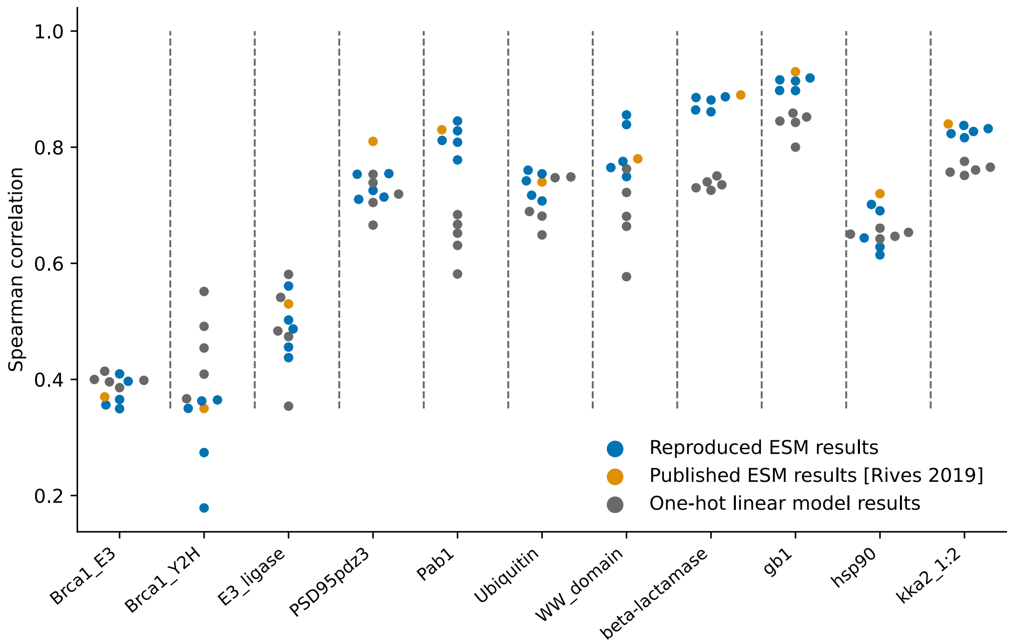


Supplementary Figure 3: **Reproducing fine-tuned Transformer results on Envision data.** When fine-tuning ESM-1b Transformer on the same Envision[2] data as used by Rives et al.[3], we reproduced the Spearman correlation performance for most of the Envision data sets on 80-20 splits. Orange is based on numbers from Supplementary Table S5 by Rives et al.[3], while blue is from our reproduced runs. For each data set, we samples 80% training data with 5 random seeds (20% of the 80% is used for validation to determine early stopping), and report the Spearman correlation on the 20% held-out data.

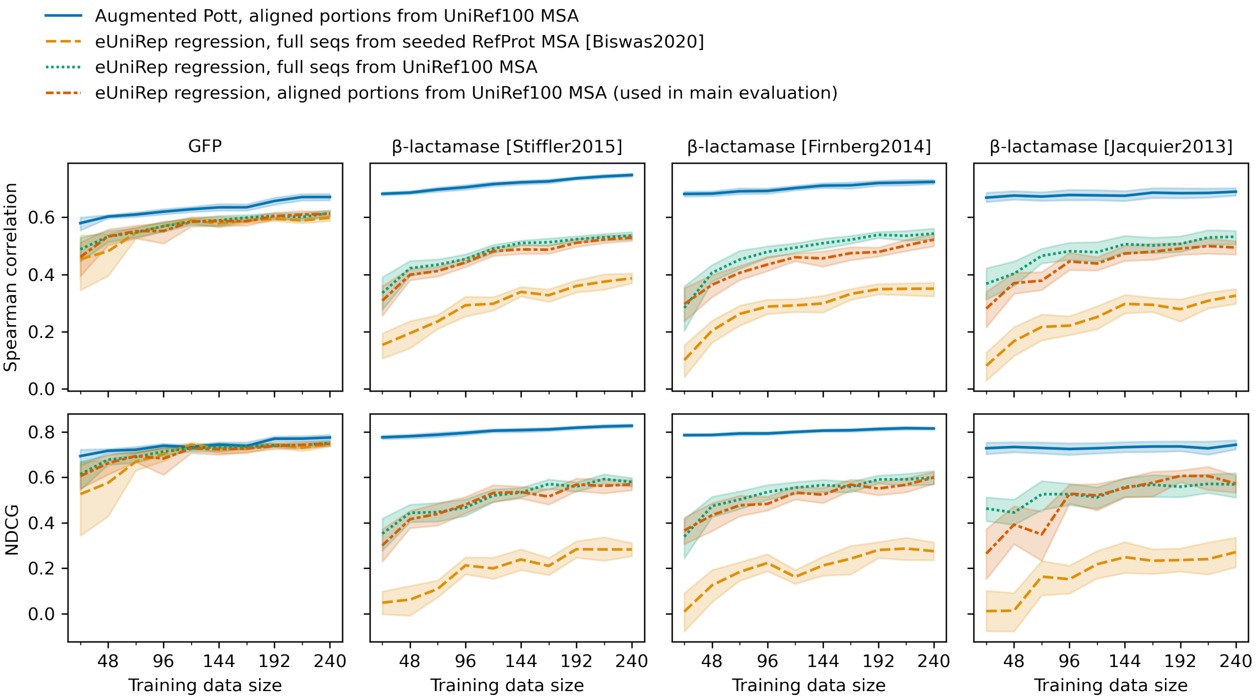


Supplementary Figure 4: **Comparison of eUniRep evo-tuning procedures**. We made two modifications in the evo-tuning procedures for eUniRep models: 1) we used the EVmutation jackhmmer parameters for consistent evolutionary data with other methods, and 2) we focused on aligned portions of sequences rather than unaligned full sequences to lower the computational burden. Here we show these two modifications do not negatively affect performance on GFP and $\beta$-lactamase—the two proteins studied by Biswas et al.[4]. The orange and green lines compare the effects of jackhmmer parameters, while the green and red lines compare the effects of using aligned portions instead of full sequences. For GFP, we used the open-sourced eUniRep model[5] as the orange line, while for $\beta$-lactamase we used Pfam PF00144 as jackhmmer seed sequences along with default jackhmmer parameters from the webserver and evo-tuned UniRep on the resulting sequences.

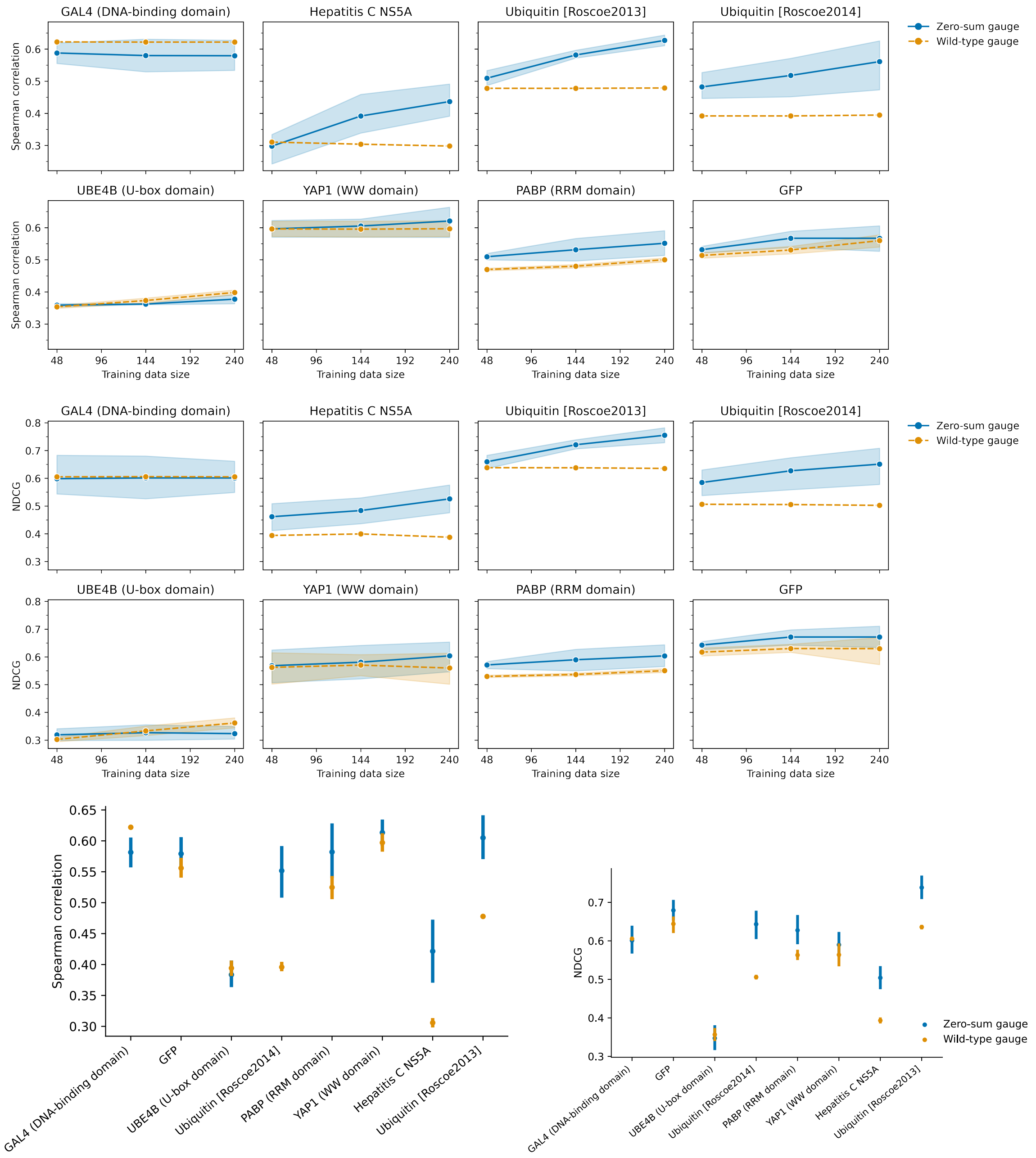


Supplementary Figure 5: **Comparison of integrated Potts models with wild-type gauge and zero-sum gauge.** Top: Spearman correlation comparisons on models trained with 48, 144, 240 training examples. Middle: NDCG comparisons on models trained with 48, 144, 240 training examples. Bottom left: Spearman correlation comparisons on 80-20 splits. Bottom right: NDCG comparisons on 80-20 splits. Overall, zero-sum gauge has better performance than wild-type gauge.

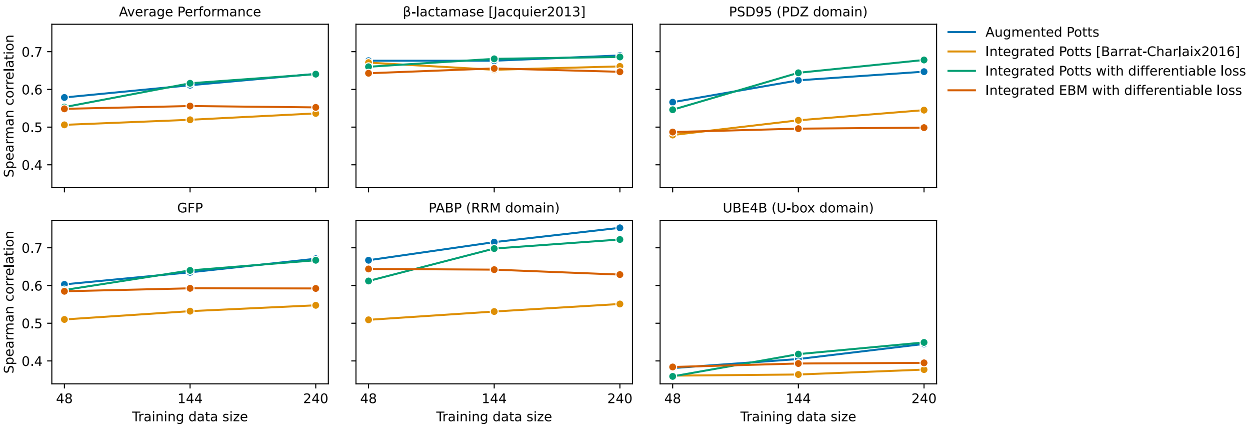


Supplementary Figure 6: **Comparison of tied-energy models and the augmented Potts model.** We evaluated three versions of tied-energy (integrated) models: 1) integrated Potts models[6] with rank-based monotonic mapping[7] between energies and fitness values; 2) integrated Potts models with a differentiable proxy of Spearman correlation as loss; 3) integrated energy-based models with neural network energy function and the same differentiable loss. As integrated models are very expensive to train, we chose to evaluate integrated models on the same $\beta$-lactamase and PSD95 data sets as Barrat-Charlaix et al.[6] and also additionally on the three data sets with higher-order mutants. In contrast to tied-energy models where evolutionary data and labelled data are fitted together, the augmented Potts model takes a two step procedure first fitting to evolutionary data and then to labelled data. While the augmented Potts model is orders of magnitude faster to compute, it still results in comparable performance with the best tied-energy model.

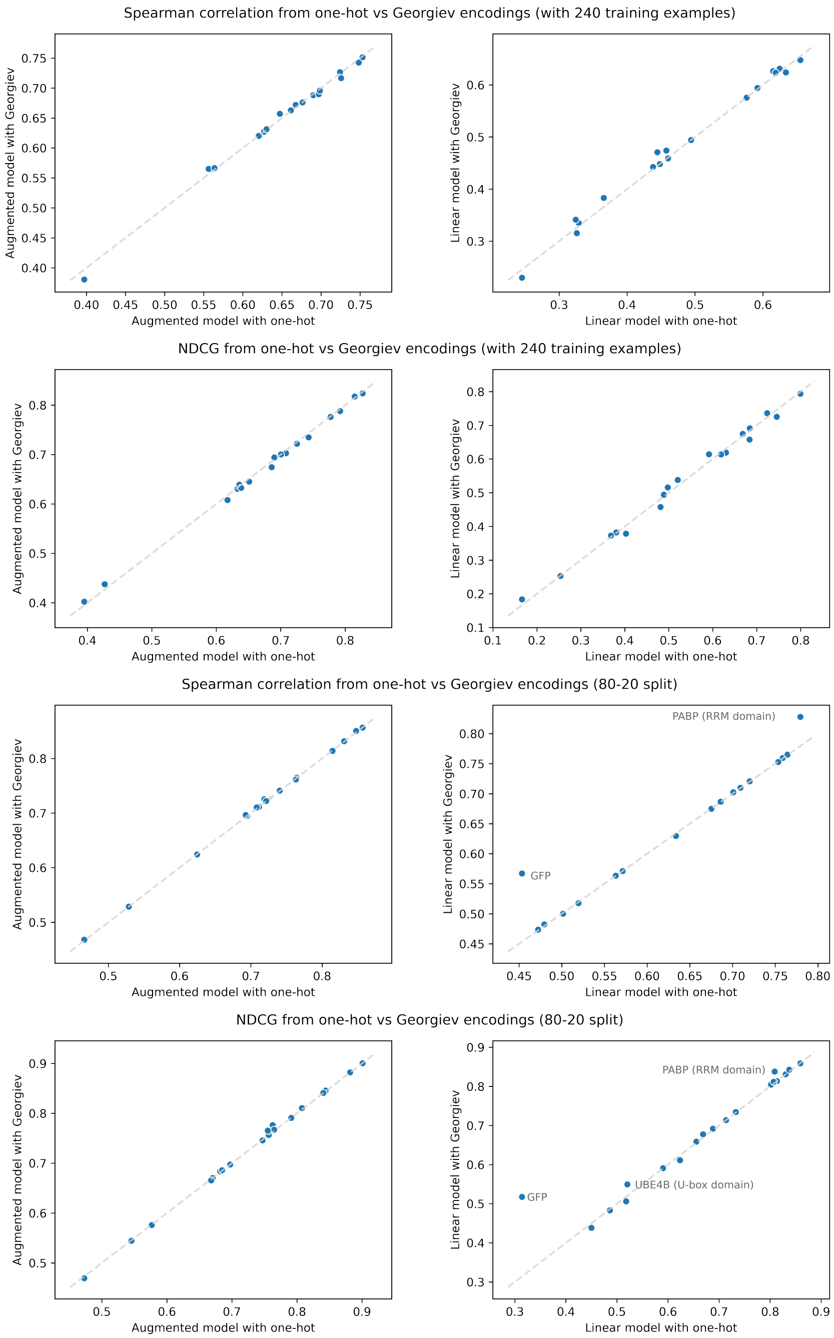


Supplementary Figure 7: **Comparison of Georgiev encoding and one-hot encoding.** We compare models with one-hot encoding to the same models with the 19-dimensional physicochemical representation of the amino acid space developed by Georgiev[8]. Each dot and error bar here represents mean and 95% confidence interval from sampling train and test data with 20 random seeds. On most single-mutant data sets, the Georgiev encoding and the amino acid one-hot encoding result in similar performance. However, on data sets with higher-order mutants when presented with sufficient training data (bottom right), the Georgiev encoding does achieve better performance than the one-hot amino acid encoding, agreeing with previous findings[9]. For the augmented Potts model, the choice of encoding does not influence performance.

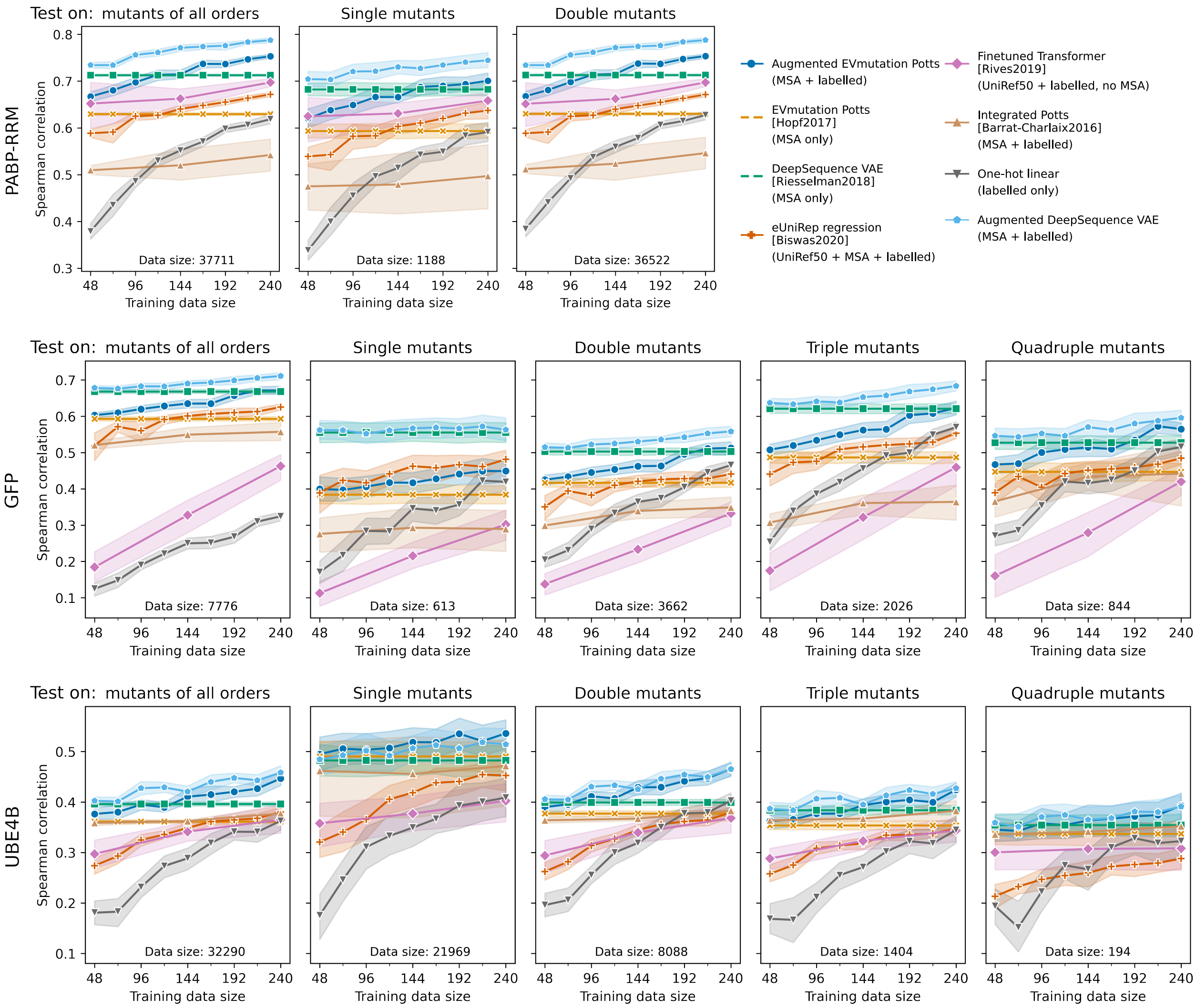


Supplementary Figure 8: **Extrapolation performance to higher-order mutants by Spearman correlation.** The “double mutants” column is a breakdown of Figure 2 top right. We test extrapolation by training on single mutants and testing on higher-order mutants on three case studies: the green fluorescent protein (GFP), the RRM domain of Poly(A)-binding protein (PABP), and the U-box domain of ubiquitination factor E4B (UBE4B). Each column measures the ranking quality among mutants that are 1, 2, 3, or 4 mutations away from the wild-type. See Supplementary Figure 9 for normalized discounted cumulative gains (NDCG) in addition to the shown Spearman correlations.

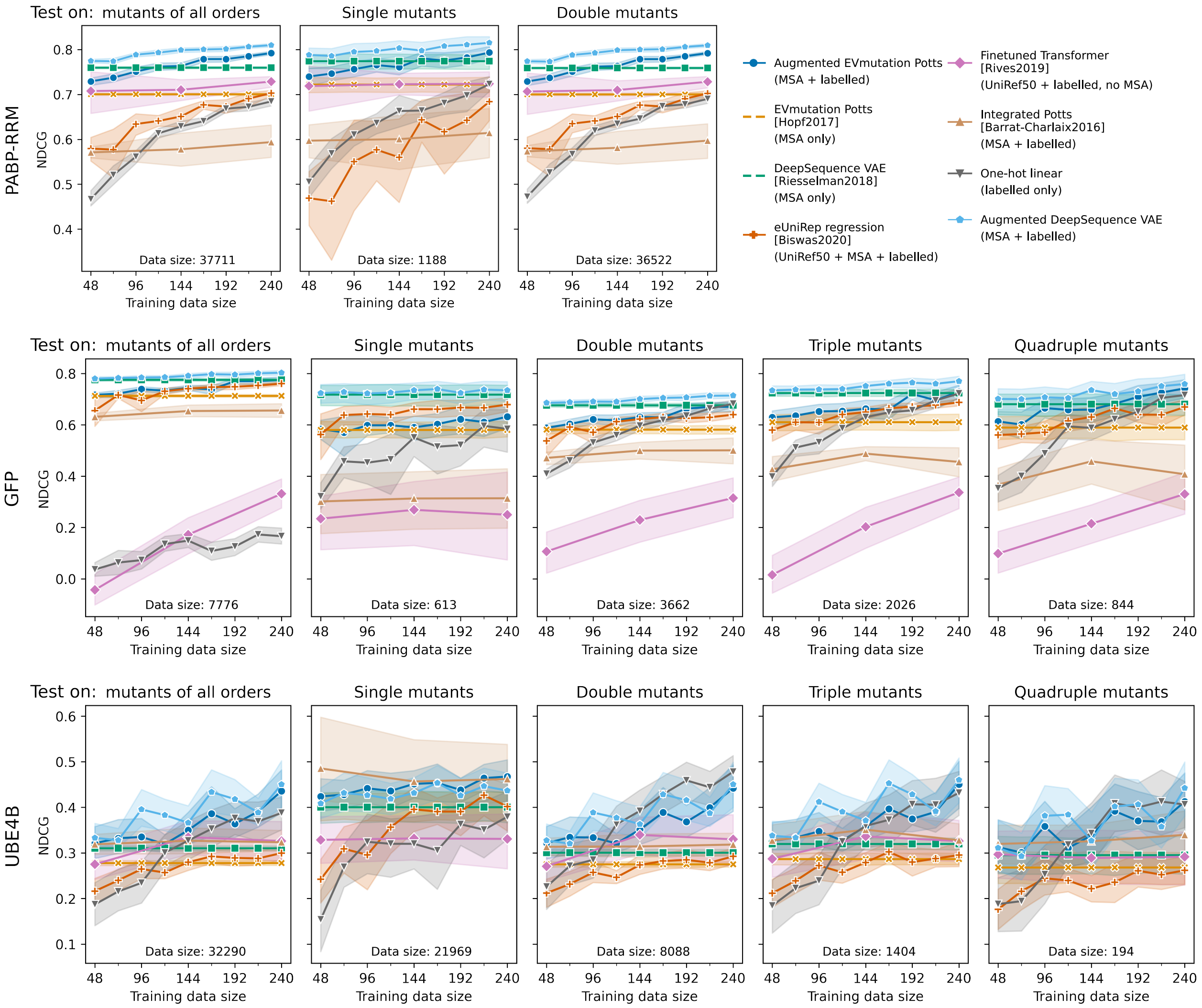


Supplementary Figure 9: **Extrapolation performance to higher-order mutants by NDCG.** The "double mutants" column is a breakdown of Figure 2 bottom right. Similar to Supplementary Figure 8, but with normalized discounted cumulative gains (NDCG) instead of Spearman correlations.

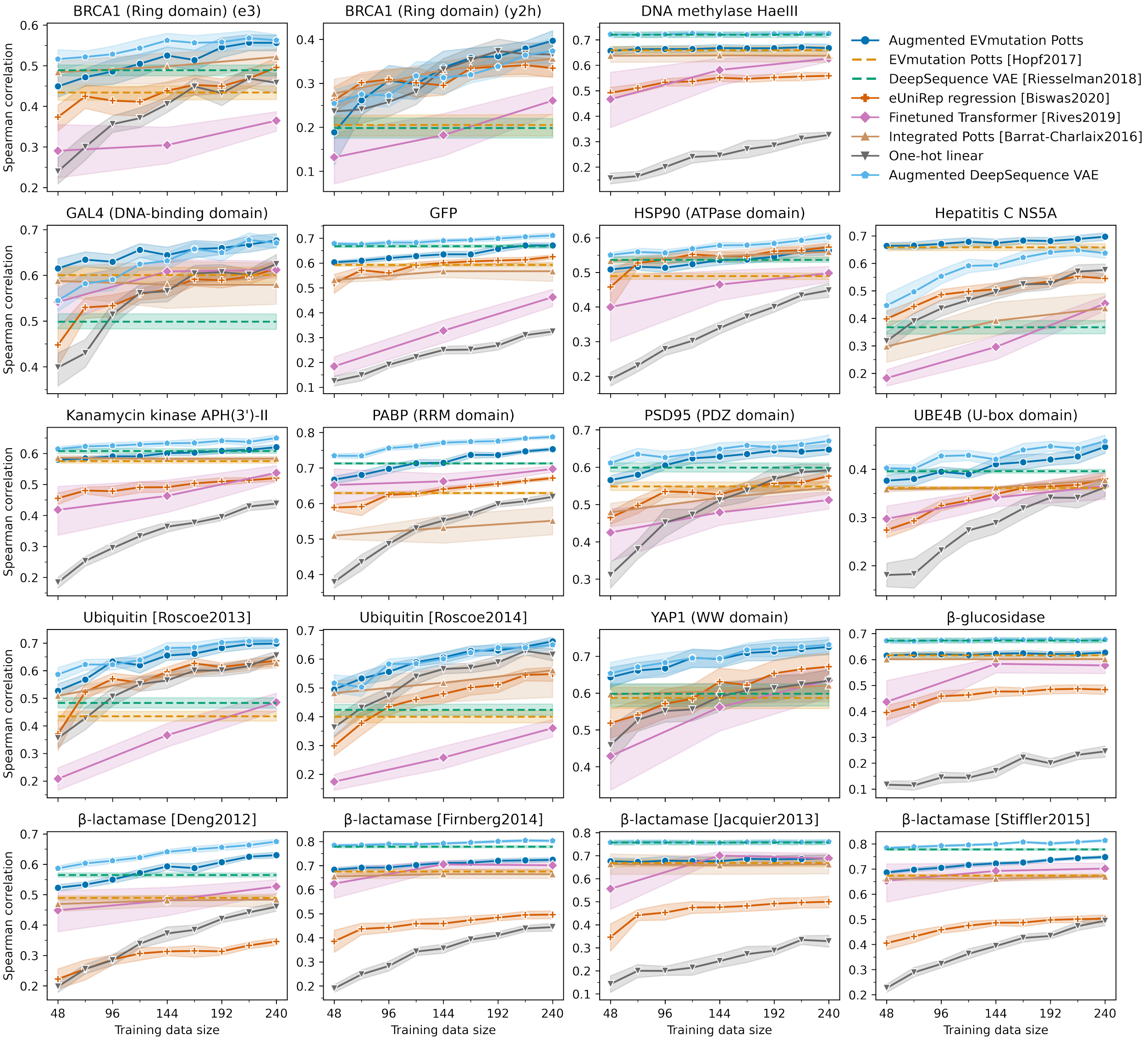


Supplementary Figure 10: **Spearman correlations with limited labelled data (breakdown of Figure 2 top left)**. See Supplementary Figure 11 for normalized discounted cumulative gains (NDCG).

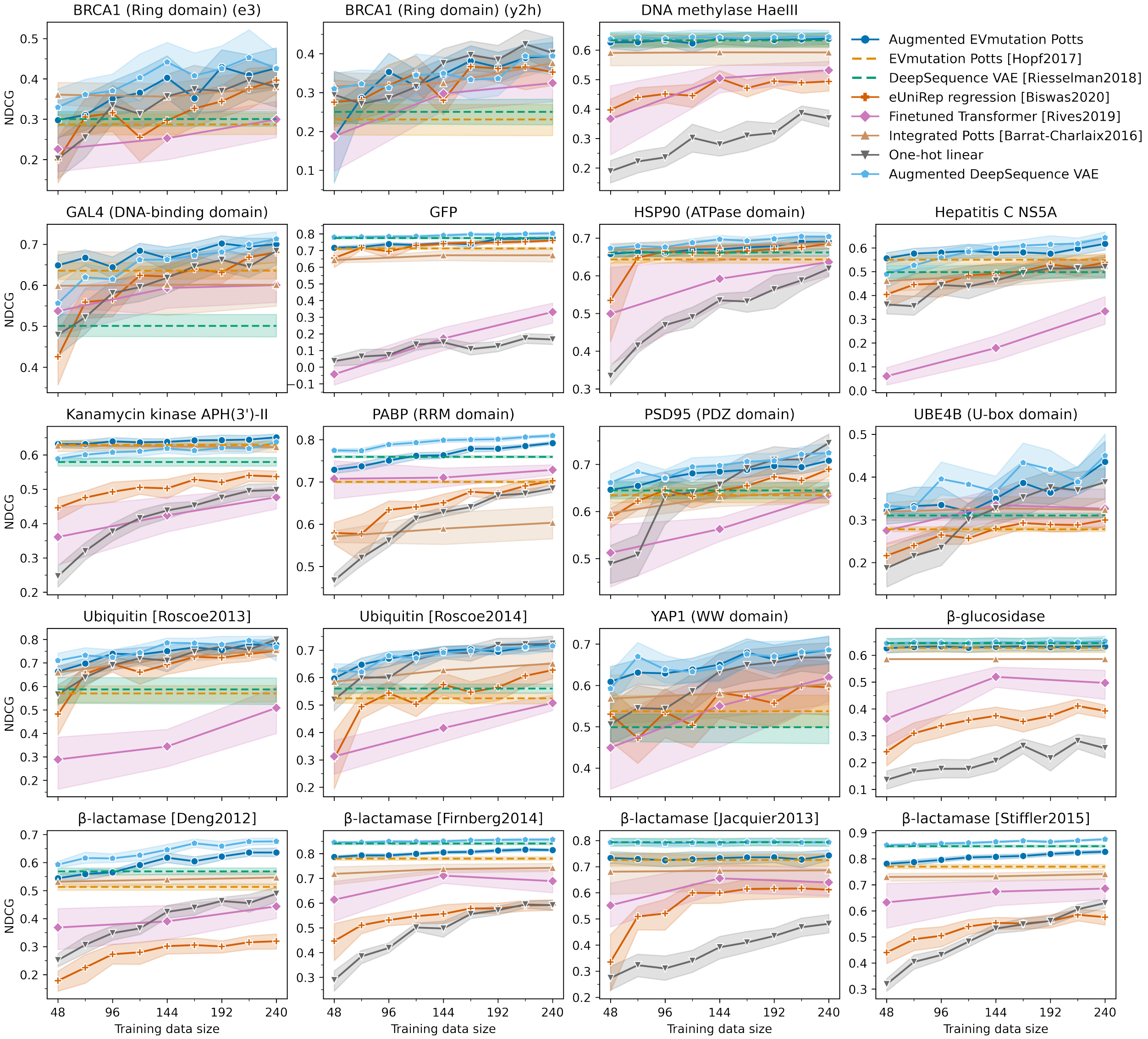


Supplementary Figure 11: **Normalized discounted cumulative gains (NDCG) with limited labelled data (breakdown of Figure 2 bottom left).** Similar to Supplementary Figure 10 but with NDCG instead of Spearman correlations.

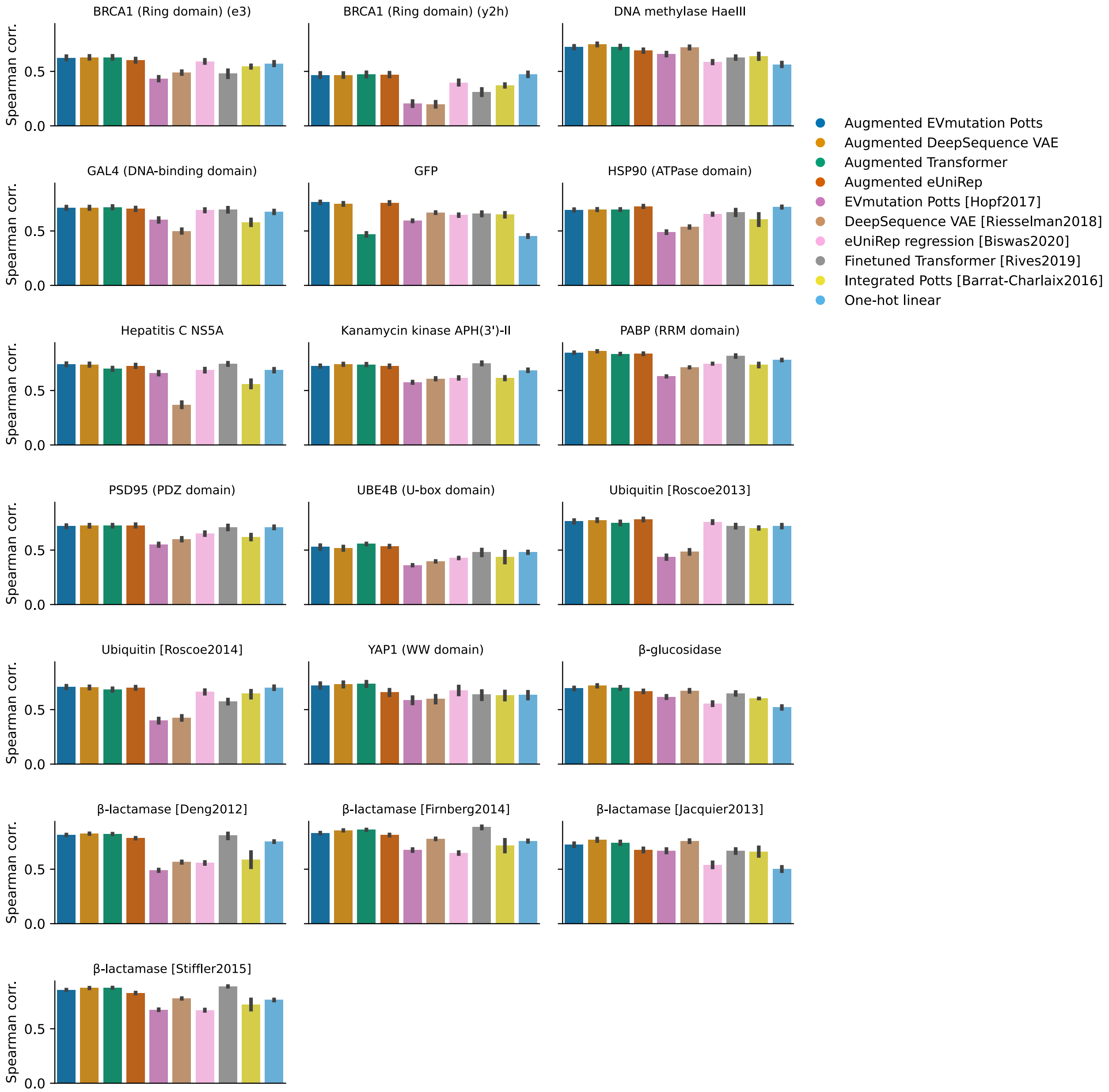


Supplementary Figure 12: **Spearman correlation on 80-20 splits (breakdown of Figure 2 top left mini-panel).** See also Supplementary Figure 13 for NDCG.

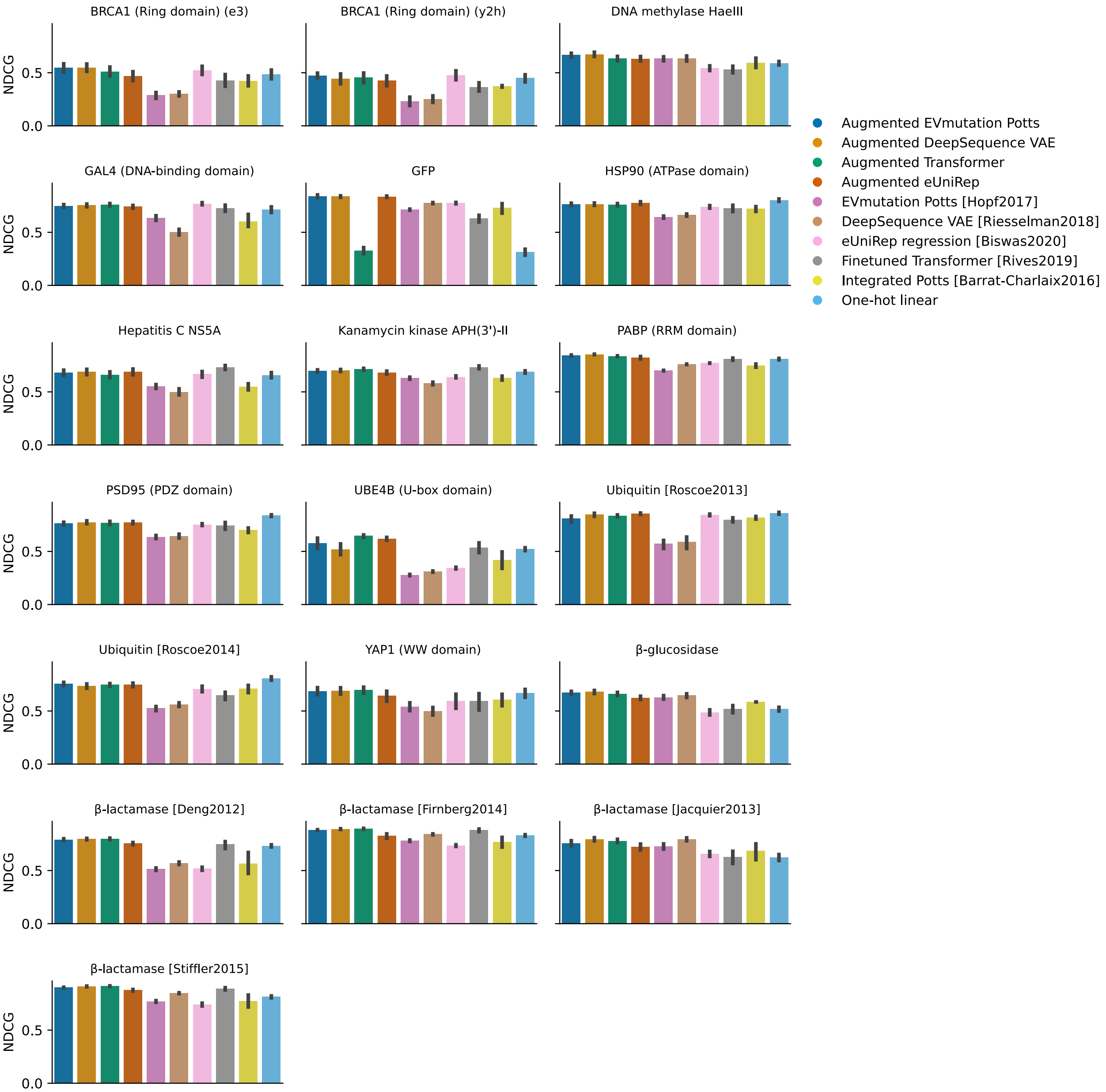


Supplementary Figure 13: **Normalized discounted cumulative gains (NDCG) on 80-20 splits (breakdown of Figure 2 bottom left mini-panel).** See also Supplementary Figure 12 for Spearman correlations.

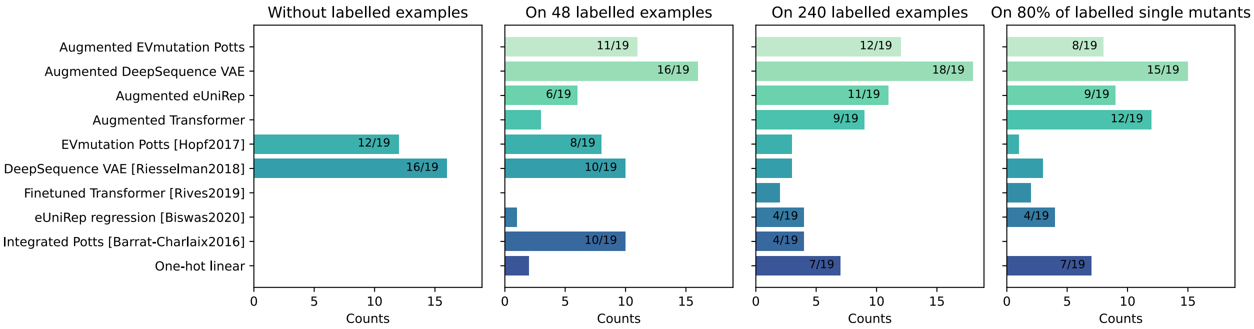


Supplementary Figure 14: **Frequency of each method achieving the highest NDCG per data set.** This figure is an alternative version of Figure 14 with normalized discounted cumulative gains (NDCG) instead of Spearman correlation for selecting the best model(s) for each data set. Similar to the observations in Figure 4, augmented models perform increasingly well on larger training data.

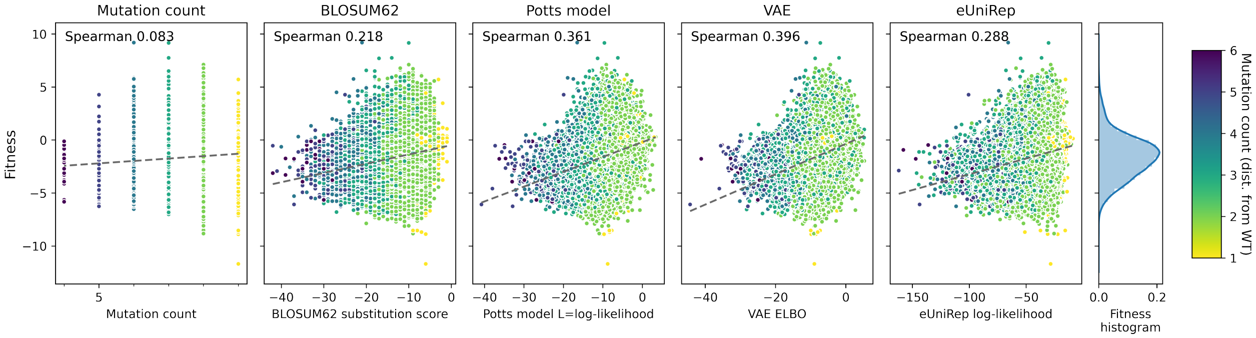


Supplementary Figure 15: **Correlations between sequence density estimations of the UBE4B U-box domain and experimentally measured fitness values.** Each dot represents a mutant sequence, with darker color indicating further distance from the wild-type. Unlike the case for GFP (Figure 5), here mutation count is not a good predictor for fitness. Moreover, none of the other methods shown here is a particularly strong predictor either.

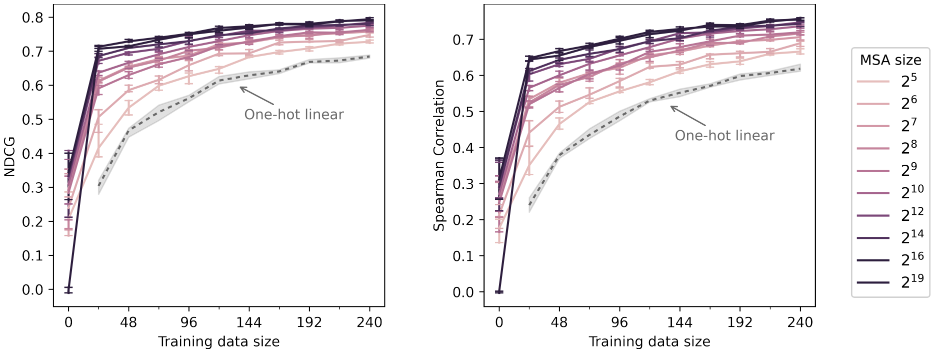


Supplementary Figure 16: **Robustness of the augmented Potts model to MSA size.** With the RRM domain of the poly(A)-binding protein as a case study, we start with the MSA obtained by jackhmmer with bit score 0.5 bits/residue, and then subsequently subsample the MSA randomly. We chose this data set for case study because it is associated with the largest MSA. The performance of the augmented Potts model degrades gradually when subsampling the MSA, but the augmented Potts model still outperforms naive linear regression even with only 32 evolutionary sequences.

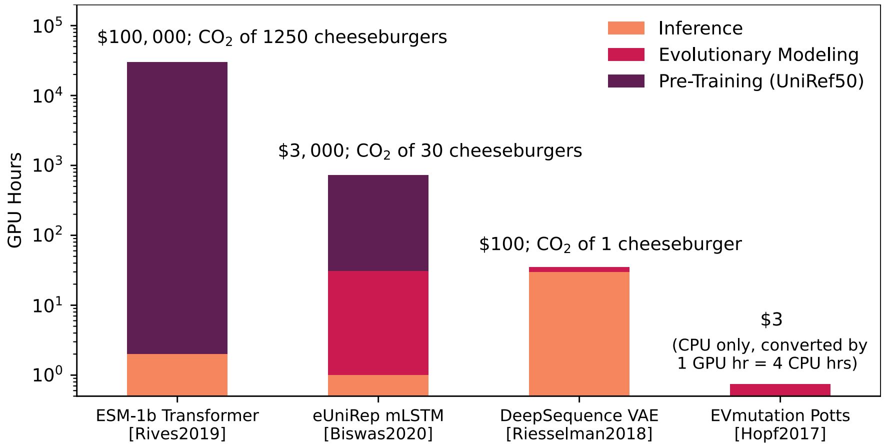


Supplementary Figure 17: **Estimation of the computing resources needed for the pre-training, evolutionary modeling, and inference of each model.** Augmented models do not require additional computing resources beyond their corresponding density models. The GPU hours for pre-training are based on the resources reported[3,5], while the hours for evolutionary modeling and inference shown here are the median running time for one data set on one Nvidia Tesla V100 GPU based on our model evaluations. GPU hours are approximately converted to costs and carbon footprints according to AWS statistics.

| Protein | UniProt ID | Measurement | # Mutations before exclusion | # Mutations after excluding positions with $\geq 30\%$ gaps | MSA size | Reference | PMID |
|---|---|---|---|---|---|---|---|
| $\beta$-glucosidase | (Sequence from dataset) | Enzyme function | 3000 (1) | 2634 (1) | 28048 | Romero et al., PNAS, 2015 | 26040002 |
| $\beta$-lactamase | BLAT_ECOLX | Growth<br>Growth<br>MIC<br>Growth | 5199 (1)<br>4997 (1)<br>990 (1)<br>4998 (1) | 4611 (1)<br>4807 (1)<br>951 (1)<br>4808 (1) | 8403 | Firnberg et al., Mol Biol Evol, 2014<br>Stiffler et al., Cell, 2015<br>Jacquier et al., PNAS 2013<br>Deng et al., JMB, 2012 | 24567513<br>25723163<br>23878237<br>23017428 |
| BRCA 1 (RING domain) | BRCA1_HUMAN | E3 ligase activity<br>BARD1 interaction | 4872 (1)<br>1748 (1) | 1382 (1)<br>1335 (1) | 25828 | Starita et al., Genetics, 2015 | 25823446 |
| PSD95 (PDZ domain) | DLG4_RAT | Peptide binding | 1578 (1) | 1578 (1) | 102410 | McLaughlin et al., Nature, 2012 | 23041932 |
| GAL4 (DNA-binding domain) | GAL4_YEAST | Growth | 1196 (1) | 1123 (1) | 17521 | Kitzmann et al., Nat Methods, 2015 | 25559584 |
| HSP90 (ATPase domain) | HSP82_YEAST | Growth | 4324 (1) | 4104 (1) | 15329 | Mishra et al., Cell Reports, 2016 | 27068472 |
| Kanamycin kinase APH(3')-II | KKA2_KLEPN | Growth | 4582 (1) | 4385 (1) | 12861 | Melnikov et al., NAR, 2014 | 24914046 |
| DNA methylase HaeIII | (Stabilized from MTH3_HAEAE) | Growth | 1778 (1, filtered) | 1634 (1) | 14115 | Rockah-Shmuel et al., PLOS Comp Bio, 2015 | 26274323 |
| Poly(A)-binding protein (RRM domain) | PABP_YEAST | Growth | 1188 (1)<br>36522 (2) | 1188 (1)<br>36522 (2) | 152041 | Melamed et al., RNA, 2013 | 24064791 |
| Hepatitis C NS5A | POLG_HCVJF | Viral replication | 1632 (1) | 1632 (1) | 8106 | Qi et al., PLOS Pathogens, 2014 | 24722365 |
| Ubiquitin | RL401_YEAST | Growth<br>E1 reactivity | 1196 (1)<br>1366 (1) | 1161 (1)<br>1295 (1) | 21448 | Roscoe et al, JMB, 2013<br>Roscoe et al, JMB, 2014 | 23376099<br>24862281 |
| UBE4B (U-box domain) | UBE4B_MOUSE | Ligase activity | 91031 (1-9 mut.) | 613 (1)<br>21969 (2)<br>8088 (3)<br>1404 (4)<br>216 (¿4) | 9172 | Starita et al., PNAS, 2013 | 23509263 |
| YAP1 (WW domain 1) | YAP1_HUMAN | Peptide binding | 363 (1) | 319 (1) | 40302 | Araya et al., PNAS, 2012 | 23035249 |
| Green Fluorescent Protein | (Sequence from paper) | Fluorescence | 51715 (1-14 mut.) | 613 (1)<br>3662 (2)<br>2026 (3)<br>844 (4)<br>630 (¿4) | 22535 | Sarkisyan et al., Nature, 2016 | 27193686 |

Supplementary Table 1: **List of all data sets.** In the "# Mutations" columns, the numbers in the brackets represent the order of mutants (*i. e.*, the Hamming distance to the wild-type sequence). For example, "12777 (2)" indicates that the data set contains 12777 double-mutant sequences.
